# Transient genomic instability drives tumorigenesis through accelerated clonal evolution

**DOI:** 10.1101/2020.11.17.387753

**Authors:** Ofer Shoshani, Bjorn Bakker, Yin Wang, Dong Hyun Kim, Marcus Maldonado, Matthew A. Demarest, Jon Artates, Ouyang Zhengyu, Adam Mark, Rene Wardenaar, Roman Sasik, Diana C.J. Spierings, Benjamin Vitre, Kathleen Fisch, Floris Foijer, Don W. Cleveland

## Abstract

Abnormal numerical and structural chromosome content is frequently found in human cancer. To test the role of aneuploidy in tumor initiation and progression, we compared tumor development in mice with chronic chromosome instability (CIN) induced by inactivation of the spindle assembly checkpoint (produced by Mad2 deficiency) and mice with transient CIN through transiently increased expression of polo-like kinase 4 (PLK4), a master regulator of centrosome number. Tumors forming under chronic CIN gradually trended toward chromosomal gains producing a specific karyotype profile that could only be partially maintained in end-stage tumors, as determined by single-cell whole genome DNA sequencing. Short term CIN from transient PLK4 induction generated significant centrosome amplification and aneuploidy resulting in formation of aggressive T cell lymphomas in mice with heterozygous inactivation of one p53 allele or accelerated tumor development in the absence of p53. Transient CIN increased the frequency of lymphoma-initiating cells (as revealed by T cell receptor sequencing) with a specific karyotype profile containing triploid chromosomes 4, 5, 14, and 15 occurring early in tumorigenesis. Overall, our evidence demonstrates that distinct CIN mechanisms drive cancers presenting specific, complex chromosomal alterations with transient CIN rapidly enhancing tumor formation by accelerating the generation of such events.

## Introduction

More than a century ago chromosome instability (CIN) leading to aneuploidy was recognized as a major hallmark in cancer^1,2^. Aneuploidy is frequently found in multiple types of cancer^3^ and is correlated with poor prognosis^4^. Although linkage of aneuploidy and tumorigenesis was proposed by Boveri in 1902^5^, the role of aneuploidy in cancer formation remains controversial. It has been proposed that increased CIN leading to aneuploid genotypes serves as a risk factor for carcinogenesis^6,7^. Indeed, reduced levels of the kinesin family motor protein CENP-E drives CIN from missegregation of individual whole chromosomes and this generates spontaneous lymphomas and lung cancers^8^. Similarly, chronic CIN from weakening of the spindle assembly checkpoint (through reduction or mutation in Bub1^9^ or from reduction in the Mps1 kinase^10^) drives tumors in mice through loss of tumor suppressor genes. Alternatively, genome doubling yielding tetraploid cells can serve as an intermediate step towards aneuploidy and cancer^11^ although tetraploidy can also protect against cell transformation^12^. Specific aneuploid backgrounds have been reported to suppress cell viability, even under oncogenic induction, thereby preventing transformation, but in cancer, aneuploidy is frequently linked to more aggressive phenotypes^13,14^. While chronic CIN at a high level can act as a tumor suppressor^15^, whether aneuploidy once developed persists in healthy cells or whether it can drive transformation or cancer initiation is not determined.

Centriole duplication is controlled by Polo-like kinase 4 (Plk4)^16,17^, a self-regulatory kinase^18^. Overexpression of Plk4 leads to centrosome amplification which can persist under p53 deficiency^18^ and drives CIN through merotelic attachment of chromosomes^19^. Indeed, centrosome amplification can be detected in pre-malignant cells in Barrett’s Esophagus and persists through malignant transformation^20^. The first direct evidence for a role of centrosome amplification in tumor formation came from flies in which increased SAK/Plk4 expression enabled tumorigenic growth of larval brain cells^21^. Chronic induction of high levels of Plk4 in mice did not increase spontaneous tumor formation even in a p53 deficient background^22^, probably due to an excessive rate of continuous CIN. However, milder chronic Plk4 overexpression has been reported to promote spontaneous tumorigenesis in wild-type^23^ and deficient p53 mice^24^. Overexpression of Plk4 during development of p53 null skin cells enhances skin cancer in adult mice^25^.

By transiently inducing Plk4 overexpression in mice we now use multiple sequencing approaches to demonstrate that transient CIN serves to enhance tumor formation by increasing tumor initiating cell frequency and by accelerating acquisition of a specific aneuploidy profile. This profile includes the gain of additional chromosomes 4, 5, 14, and 15, a tumor karyotype we demonstrate to develop early during tumorigenesis under transient CIN, and also in end-stage tumors forming under chronic CIN from inactivation of the spindle assembly checkpoint (SAC). Transcriptomic analysis revealed that this aneuploidy profile drives a gene expression program that is also frequently observed in multiple human cancers. We conclude that transient CIN is a powerful mechanism driving tumors with a specific aneuploidy profile.

## Results

### Aneuploidy selection in tumors under chronic CIN driven by deletion of Mad2

We initially used single-cell whole genome DNA sequencing (scWGS) to examine aneuploidy evolution during the development of tumors formed after silencing the SAC by selective inactivation in the thymus of both alleles encoding Mad2, thereby driving chronic CIN (Supplementary Figure 1a-b). Thymic tumors of different sizes were collected after Cre-recombinase (encoded by Lck promoted Cre transgene) mediated homozygous inactivation in T lymphocytes of “floxed” Mad2 and p53 genes in mice. Selective Mad2 deletion (in Lck-Cre+; Mad2f/f; p53f/f mice) occurs ~6 weeks of age when Lck expression initiates in T cells^26^.

We previously showed that Lck-Cre; Mad2f/f; p53f/f mice succumb to thymic lymphomas within ~4 months with recurrent aneusomies involving chromosomes 4, 5, 14, 15 and 17^26^. To determine when in tumorigenesis these clonal karyotypes emerged, thymuses were harvested from mice aged between 8 and 16 weeks representing a spectrum of tumor development from early (100-500mg) to advanced (>500mg) thymic lymphomas (Figure 1a). Each was homogenized and single-cell suspensions were used for single cell whole-genome sequencing and bulk homogenates were used for transcriptome analyses (RNA sequencing or RNA-seq) (Figure 1a). Initially, to determine the influence of SAC inactivation on potential aneuploidy development in wild-type p53 mice, analyses from two Mad2f/f; Lck-Cre+ thymuses from 8-week old mice revealed 48% and 46% aneuploidy (with aneuploidy scored by either gains or losses of at least one chromosome). No selection for specific chromosome copy number changes was identified (Figure 1b and Supplementary Figure 1c).

**Figure 1.**
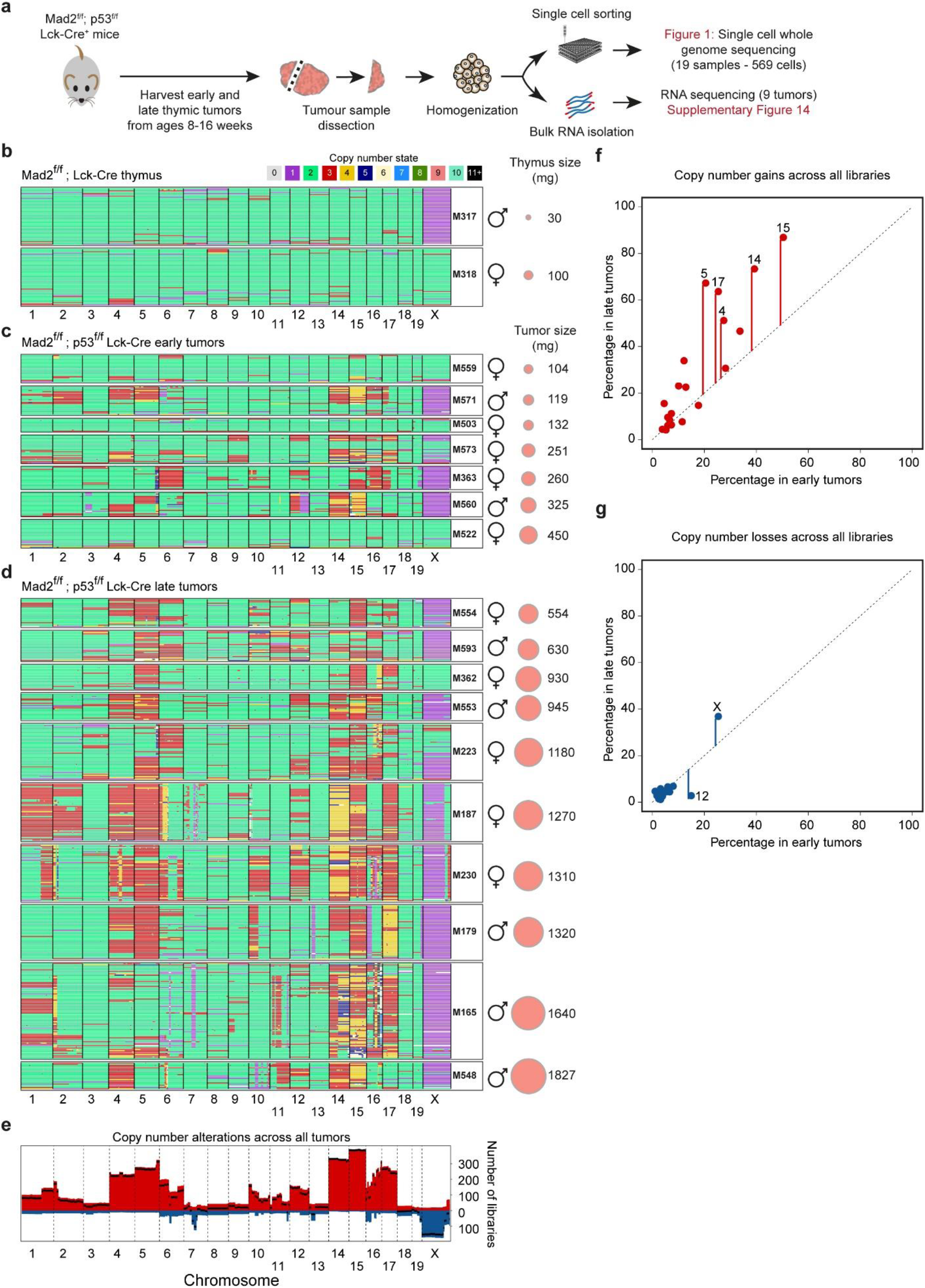
Aneuploidy evolution in thymic lymphomas under chronic CIN driven by deletion of Mad2. (**a**) Overview of thymic tumors collection from Lck-Cre+; Mad2f/f; p53f/f mice sampled at 8-16 weeks of age. (**b-d**) Heatmaps showing DNA copy number using single cell whole-genome sequencing of tumor cells collected from (**b**) Lck-Cre+; Mad2f/f mice (**c**) Lck-Cre+; Mad2f/f; p53f/f mice with developing (early) tumors and (**d**) Lck-Cre+; Mad2f/f; p53f/f mice with terminal (late) tumors. Genomic position in order from chromosome 1 to X are in the x-axis and individual cells are in the y-axis. Colors indicate the copy number state as determined by AneuFinder. Data of samples M187 and M179 was previously reported in Foijer et al. elife, 2017. Sample M165 is a new sequencing experiment of a sample also shown in Foijer et al. elife, 2017. (e) Genome-wide overview of cumulative copy number (1 Mb bins) gains (red) and losses (blue) across all thymic lymphomas presented in panels **c-d**. Black line presents the net change; difference between number of libraries with a copy number gain and the number of libraries with a copy number loss. (**f-g**) Comparison of copy number gains (**f**) and losses (**g**) between early and late tumors. Each dot represents one chromosome.

Genomic examination of developing (early tumors, Figure 1c) or advanced (late tumors, Figure 1d) thymic lymphomas revealed large chromosome changes, with an overwhelming majority of tumor cells presenting chromosome gains accompanied by few or no losses. Gains of chromosomes 4, 5, 14, 15 and 17 (Figure 1e) became more frequent in advanced tumors (Figure 1f-g). Analysis of tumor genome heterogeneity and copy number (CN) changes scores (Supplementary Figure 1d, see Methods for detailed definitions) revealed increased CN changes, but not heterogeneity, correlated with tumor progression. Some of the developing lymphomas, most notably M503, M522 and M559, showed aneuploidy landscapes similar to the Mad2f/f; Lck-Cre+ thymuses (Compare Figures 1b and 1c), with similar CN scores, heterogeneity, and percentage of aneuploid cells (Supplementary Figure 1c-d), indicating that selection for chromosome aberrations had not (yet) occurred. The lack of selection for chromosome aberrations is illustrated clearly in M522 whose thymus weight was near 500 mg, indicative of tumor development, although most single cells sequences were euploid with others only showing random aneusomies. In contrast, other developing lymphomas showed signs of aneuploidy karyotype selection indicated by preferential chromosome copy number changes that became more apparent in endpoint lymphomas (Figure 1f-g). An increase in the number of copy number transitions indicative of structural damage within chromosomes was also observed in advanced tumors (Supplementary Figure 1d). Thus, chronic CIN from SAC inactivation yields thymic tumors that have gradually acquired an aneuploidy profile with chromosome copy number increases in chromosomes 4, 5, 14, 15, and 17.

### Transient PLK4 overexpression leads to transient chromosome instability

We next determined how transient chromosome instability affected tumorigenesis by generating a mouse with an inducible Plk4 gene in the background of different p53 genotypes. This was achieved by two rounds of breeding (Figure 2a) to generate mice (to be referred to as **PRG5** mice) in a congenic background that carried 1) a doxycycline-regulated **<u>P</u>**LK4-EYFP gene^23^, 2) a gene encoding the **<u>R</u>**everse tetracycline transactivator (rtTA) whose expression permitted doxycycline-inducible expression of PLK4 (Supplementary Figure 2a), 3) a gene encoding centrin-**<u>G</u>**FP^27^, and 4) in which neither, one, or both p**<u>5</u>**3 alleles were inactivated^28^.

**Figure 2.**
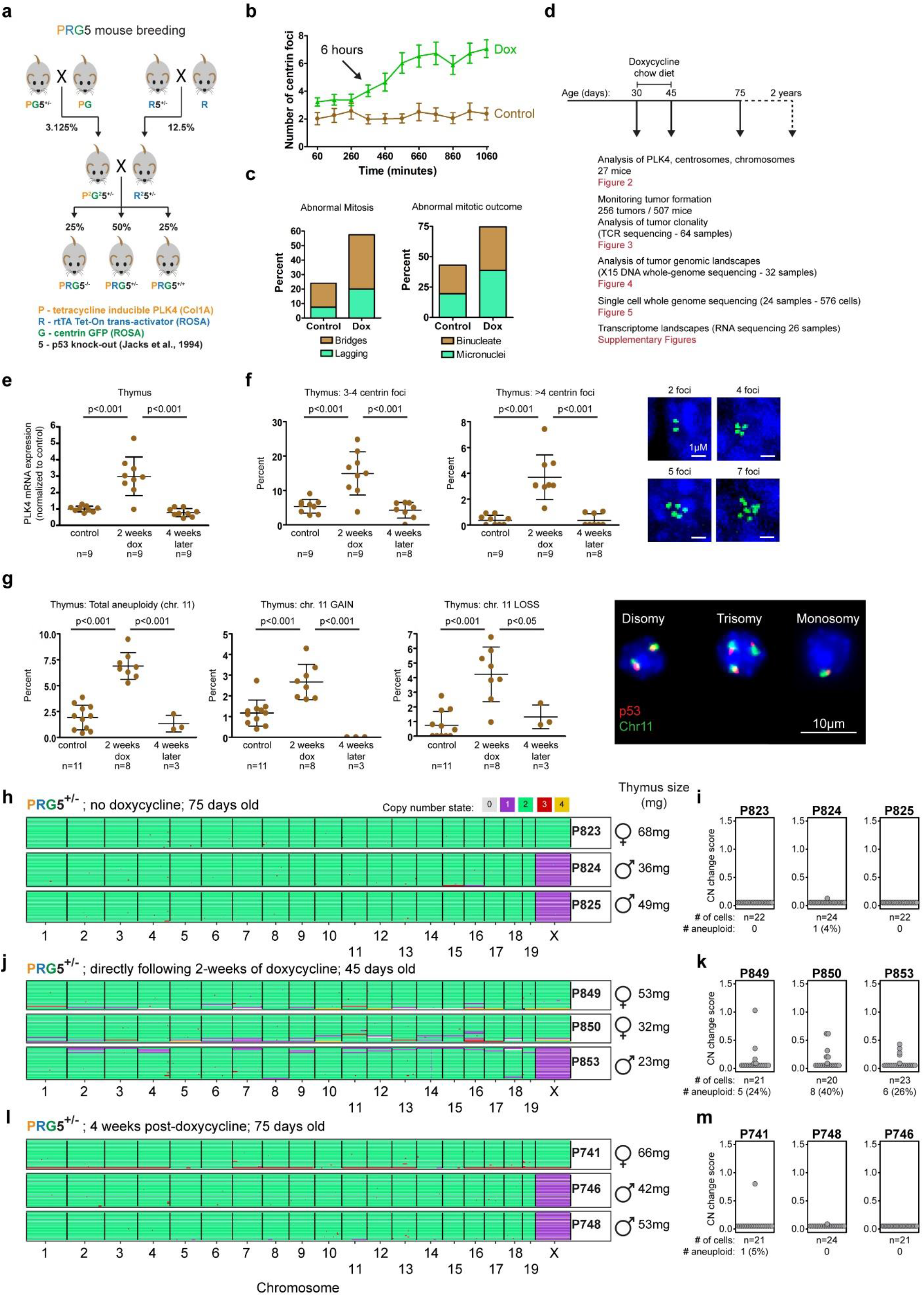
Transient CIN in mice through induced Plk4 overexpression drives transient aneuploidy. (**a**) Breeding strategy used to obtain doxycycline-inducible Plk4 mice with centrin-GFP and under different backgrounds of p53 (PRG5 mice). (**b-c**) Measurements of centrin-GFP foci (b, starting one hour following Plk4 induction, mean ± SD) and mitotic abnormalities (c, as observed starting 8 hours after Plk4 induction) in p53+/- PRG5 mouse embryonic fibroblasts using live cell imaging. (**d**) Overview of the experimental design using PRG5 mice. (**e-g**) Plk4 mRNA levels (**e**), measurement of centrin-GFP foci (**f**), and percent aneuploidy for chromosome 11 (using interphase DNA-FISH, **g**) in thymuses from PRG5 mice before (control) immediately after (2 weeks dox) and one month after (4 weeks later) doxycycline administration. Mean ± SD of indicated mice per group are presented. *p-values determined using one-way ANOVA with Tukey’s Multiple Comparison Test. (**h, j, l**) Heatmaps showing DNA copy number using single cell whole-genome sequencing of cells collected from p53+/- PRG5 mice (**h**) before, (**j**) immediately after, and (**l**) one month after Plk4 induction. (**I, k, m**) Analysis of copy number changes of the samples on the left (panels **h, j,** and **l**). See methods for details about calculation of the CN (copy number) change score. Statistics below the dot plots indicate the total number and percentage of cells with at least one whole chromosome gain or loss.

Doxycycline-dependent PLK4 induction was initially validated independent of p53 status in mouse embryonic fibroblasts (MEFs) derived from PRG5 mice. Increased (4-fold) PLK4 RNA levels and centrosome numbers (up to 6-fold) were found within two days of PLK4 induction (Supplementary Figure 2b-c). Live cell imaging revealed that centrosome amplification initiated within 6 hours after PLK4 induction (Figure 2b), producing >50% abnormal defective mitoses and binucleated daughter cells or cells with micronuclei (Figure 2c). PRG5 mice fed for two weeks with doxycycline-containing food (Figure 2d), had increased PLK4 RNA levels in the thymus (3-fold), spleen (2.5-fold), liver (2-fold), colon (20-fold), and skin (20-fold), but not in lung or kidney (Figure 2e and Supplementary Figure 2d). Plk4 induction was transient, as PLK4 RNA levels returned to basal levels within one month after discontinuing doxycycline.

Increased PLK4 RNAs in thymic cells produced transient centrosome amplification (Figure 2f and Supplementary Figure 2e) and CIN, leading to aneuploidy just after doxycycline removal with both gain and loss of chromosome 11 copy numbers in thymocytes (3-fold increase in aneuploidy, Figure 2g) as determined using interphase DNA fluorescent in-situ hybridization (FISH), but not in splenocytes (where there was only mild PLK4 RNA induction - Supplementary Figure 2d,f). Aneuploidy levels determined on a whole genome scale using scWGS (Figure 2h-m) revealed a transient increase in aneuploidy from near zero (0-4% before Plk4 induction) to 24-40% after transient (two-week Plk4 induction) CIN, which returned to near zero within one month after doxycycline withdrawal. Importantly, transient CIN produced heterogenous aneuploidy, with some single cells gaining/losing one chromosome and some gaining/losing up to 9 chromosomes (Figure 2h-m).

### Transient CIN drives thymic lymphoma initiation and acceleration

Survival and tumor formation were monitored for more than two years in PRG5 mice following two-week CIN induction beginning at 4 weeks of age (Figure 2d). In mice with wild-type p53, there was no decreased survival (median survival 837 days and 805 days in control and treated mice, respectively - Figure 3a) and no increase in tumor formation (17/42 and 13/49 control and treated mice with tumors, respectively - Figure 3b and Supplementary Figure 3a) as previously observed^10,26^. Tumor spectrum profile in aged PRG5 p53+/+ mice (Figure 3c) was also not affected by PLK4 induction, with a majority of tumors forming in spleen (~40%), the digestive system (colorectal and intestinal tumors ~20%), and liver (~17%).

**Figure 3.**
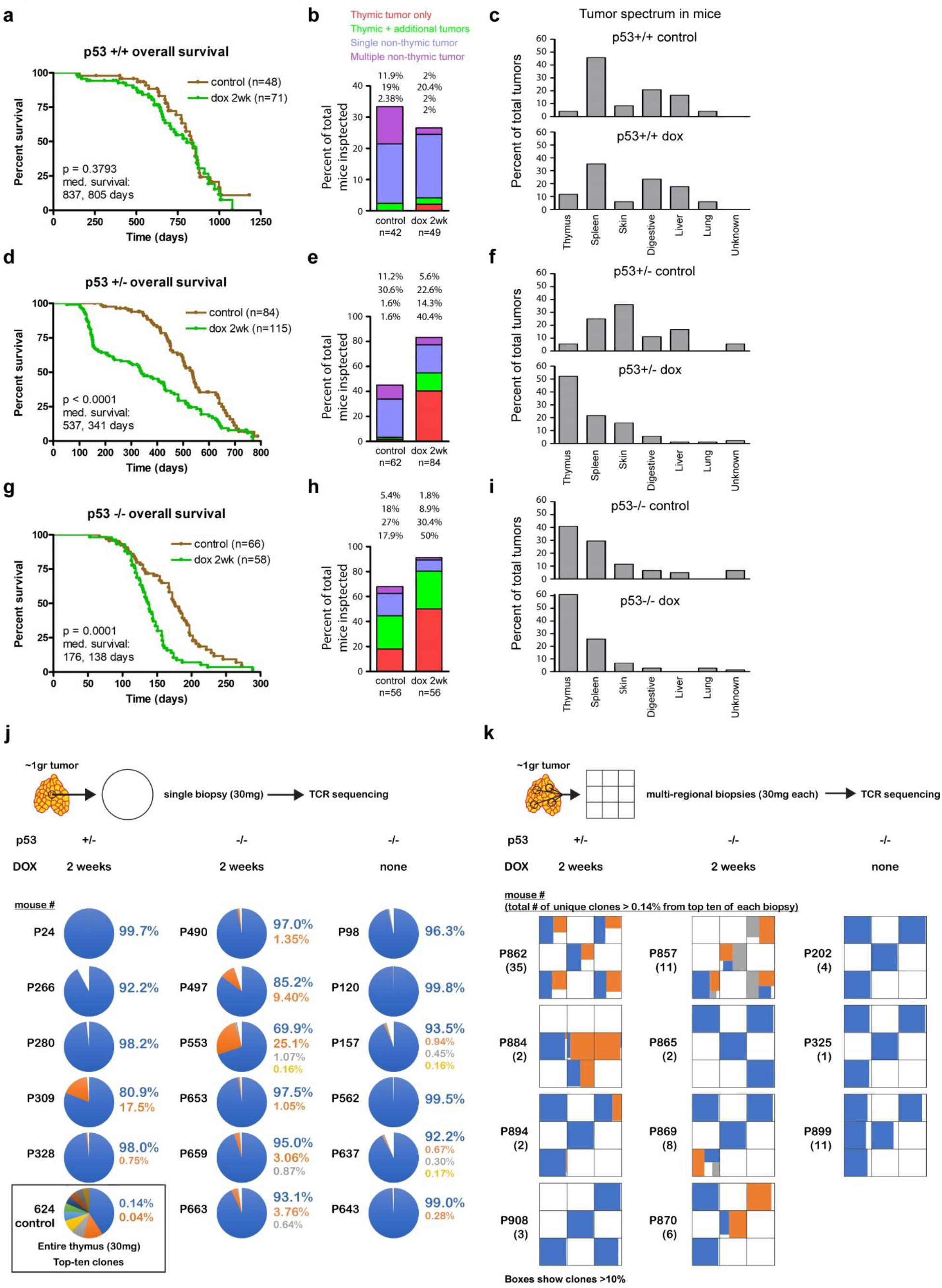
Transient CIN drives thymic lymphoma initiation in p53 deficient mice. (**a, d, g**) Survival (Kaplan-Meier) plots of PRG5 mice treated with doxycycline for two weeks at the age of 30 days with (**a**) wild-type p53, (**d**) heterozygous p53, and (**g**) p53 knock-out backgrounds. Comparison of indicated number of mice done using log rank test. (**b, e, h**) Thymic and non-thymic tumor frequencies of indicated number in PRG5 mice treated with doxycycline for two weeks at the age of 30 days with (**b**) wild-type p53, (**e**) heterozygous p53, and (**h**) p53 knock-out backgrounds. (**c, f, i**) Distribution of tumor types from indicated number of tumors in PRG5 mice treated with doxycycline for two weeks at the age of 30 days with (**c**) wild-type p53, (**f**) heterozygous p53, and (**i**) p53 knock-out backgrounds. (**j-k**) Top ten T cell receptor frequencies (indicative of T cell clones) in thymic T cell lymphomas from PRG5 mice in (**j**) single biopsies and (**k**) multi-regional biopsies as determined using T cell receptor sequencing. A control thymic sample (mouse #624) is presented in **j**.

Tumors in PRG5 mice heterozygous for p53 (overall frequency of 45.1%) were identified as early as 7 months of age, but thymic tumors were rare (3.2%). In contrast, two weeks of transient CIN led to a doubling in tumor frequency (83.3%) and reduced survival (median 341 vs. 537 days - Figure 3d), with very early onset tumor formation (starting at 100 days of age - Supplementary Figure 3b-c), the majority (54.8%) of which were thymic tumors (Figure 3e) and a clearly altered tumor spectrum profile (Figure 3f). Halving the period of CIN to only one week produced no effect on survival (median survival – 556 days vs. 507 days of uninduced mice) and only 1/10 mice presented a thymic lymphoma (Supplementary Figure 3d-f). In contrast, a longer period of CIN (PLK4 induction with doxycycline for 4 weeks) shortened survival time further (median survival – 158 days vs. the 341 days in mice with 2 weeks of CIN), with a comparable frequency of thymic lymphomas (54% vs. 55% in mice with 2-weeks of CIN). Induction of CIN at a later age (100 days instead of 30 days old mice) yielded indistinguishable survival and tumor formation frequencies (Supplementary Figure 3g-i).

p53 knockout mice succumb to tumor formation within the first year of life (median survival of 176 days and 67.8% of examined mice had detectable tumors - Figure 3g-h), with a large proportion with thymic tumors (44.6% - Figure 3h). Following two weeks of doxycycline-induced CIN, PRG5 mice with p53 knockout developed a significantly higher number of tumors (91% - Figure 3h) of which 80.4% were of thymic origin. Survival was further decreased (median survival - 139 days - Figure 3g) relative to p53-/- mice due to the increased incidence of thymic lymphomas from which mice died at an earlier age (median survival of mice with thymic lymphomas - 134 and 154 days in doxycycline treated and control mice, respectively, Supplementary Figure 3j-l), similar to observations in Lck-Cre; Mad2f/f; p53f/f mice^26^. Tumor spectrum profiles were similar with the exception of increased thymic tumor proportion (Figure 3i). Finally, survival of PRG5 mouse cohorts (either p53 knockout or heterozygote) with high frequency of thymic lymphomas was comparable (Supplementary Figure 3m), suggesting that spontaneous tumor initiation in p53-/- mice occurs after weaning. In all cases examined thymic tumors were identified as lymphomas, of which many appeared invasive and metastatic (Supplementary Figure 3n, and Supplementary table 1).

### Transient CIN drives p53 loss and Myc overexpression

RNA expression profiles of tumors and normal thymuses were determined to examine mechanism(s) underlying the conversion of normal tissue into spontaneous or doxycycline-induced thymic lymphomas. A general profile distinct from control normal thymus samples was identified for lymphomas (1325 differentially expressed genes p<0.05, Supplementary Figure 4a and Supplementary Table 2), with significant alterations in cell cycle and metabolic pathways (Supplementary Figures 4b and 5). Lymphomas from transient CIN in PRG5 p53+/- mice had low expression of Trp53 (indicative of loss of the wild-type p53 allele), low expression of p53 target genes including Bcl2l1, and activation of p19 (Supplementary Figure 6a). Indeed, expression of p53 exons 2-6 (the region deleted in the p53 null allele) was completely lost (Supplementary Figure 6b) ^28^. Whole genome DNA sequencing (X15 coverage) confirmed the loss of wild type *TRP53* gene exons 2-6 without evidence of other numerical or structural alterations in chromosome 11 (Supplementary Figure 6c-d) indicating whole-chromosome missegregation is the underlying mechanism.

Several common features were identified as hallmarks of advanced thymic lymphomas in PRG5 mice (p53-/- and p53-/- or p53+/- after transient CIN), including increased centrosome numbers (Supplementary Figure 7a) and overexpression of Myc RNA (Supplementary Figure 7b) and protein (Supplementary Figure 7c), potentially due to acquisition of additional chromosome 15 copies, an event more frequently observed in tumors from transient CIN (Supplementary Figure 7d).

### Transient CIN generates tumor initiating cells

We hypothesized that acceleration of tumor formation following transient CIN might occur due to CIN enhancing generation of tumor initiating cells. To test this, we collected small biopsies (~2-3% of the tumor) from multiple tumors, as well as multi-focal biopsies from additional tumors and sequenced the beta chain of the T cell receptors (TCRβ) found in the tumor T cells (Figure 3j-k). Single biopsy analysis revealed that tumors forming after two-weeks of induced CIN had increased clone frequencies, with 1/5 tumors in p53 +/- mice and 6/6 tumors in p53-/- mice having more than one dominant clone (>1% frequency). In contrast, no tumors (0/6) from p53-/- mice without transient CIN had more than one dominant clone (Figure 3j and Supplementary Table 3). Transcriptomes generated by RNA sequencing confirmed the specifically increased expression of individual variable TCRβ genes in tumors from PRG5 mice (Supplementary Figure 8a), and also revealed increased clonal frequency in tumors from mice with Mad2 deletion (Supplementary Figure 8b).

Tumor clonality was more directly addressed in PRG5 mice by a collection of multi-regional biopsies representing different spatial regions of a tumor. TCRβ sequencing revealed 3/4 of CIN induced p53+/- and 3/4 of CIN induced p53-/- derived tumors in PRG5 mice contained more than one dominant clone, whereas none of the spontaneous tumors from p53-/- mice contained more than one dominant clone (0/3) (Figure 3k and Supplementary Table 3). Tumors containing more than one TCRβ clone showed spatial separation of the different TCRβ receptors, consistent with independent tumorigenic events forming in parallel within the thymus (Figure 3k).

### Transient CIN accelerates clonal evolution of thymic lymphoma

Since CIN drives random genomic reshuffling and generates higher tumor initiating cells (Figure 3j-k), transient CIN could be expected to generate tumors with diverse gene expression profiles. However, this was not the case: transcriptomes of independent tumors formed in PRG5 mice following transient CIN were found to share a similar gene expression profile whereas spontaneous tumors in p53-/- mice were more heterogenous (Supplementary Figure 9a-b). Whole genome sequencing of independent tumors revealed common chromosome gains in tumors forming after transient CIN (Figure 4 and Supplementary Figure 9c), with increases in chromosomes 4, 5, 14, and 15. RNA expression levels confirmed the apparent changes in chromosome copy numbers (Supplementary Figure 9d).

**Figure 4.**
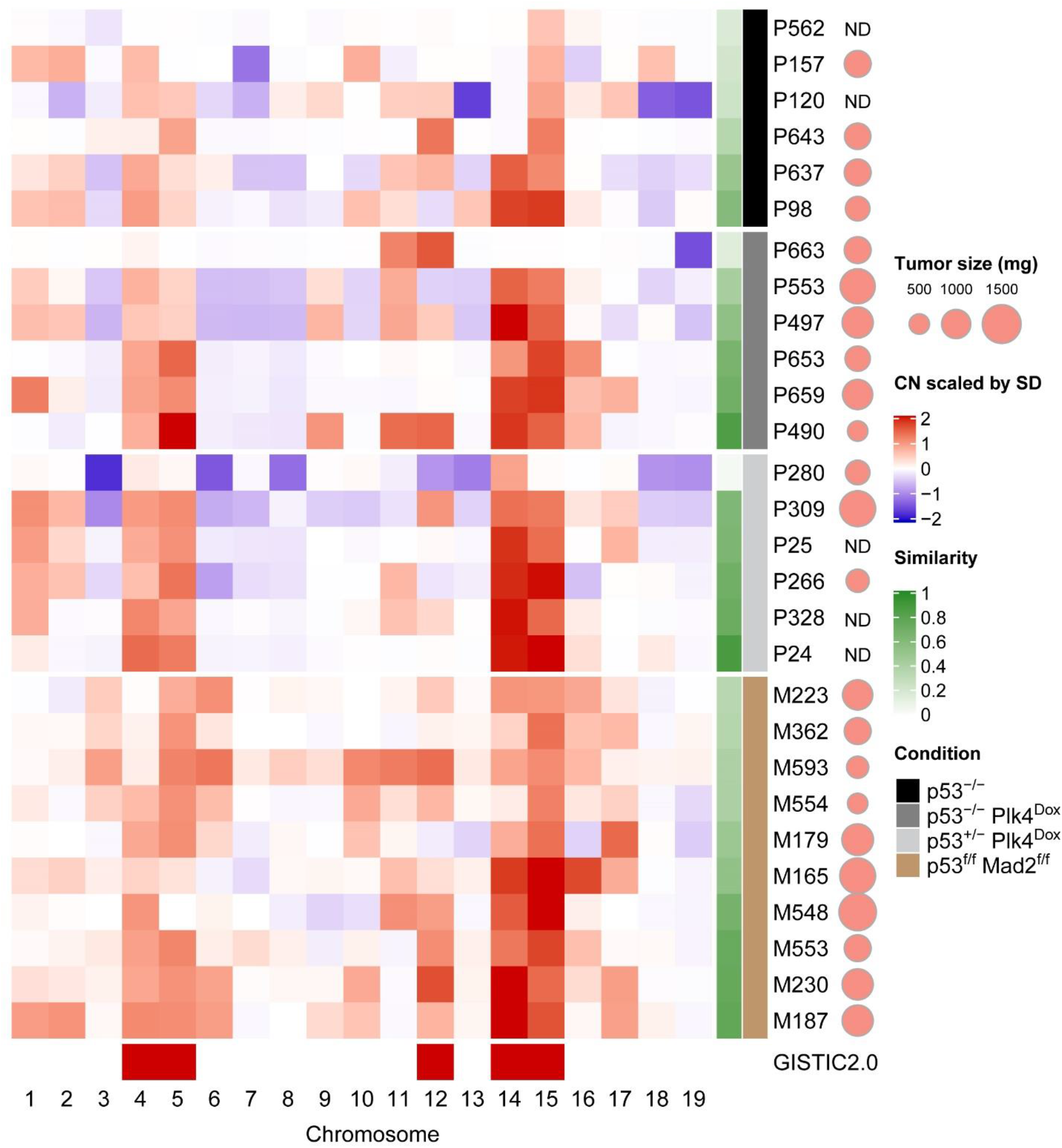
CIN accelerates the selection for a specific aneuploidy profile. Heatmap showing averaged chromosome DNA copy number changes in late tumors (>500mg) from non-induced p53-/- PRG5 mice (black), two-weeks doxycycline treated (at the age of 30 days) p53-/- PRG5 mice (dark grey) and p53+/- PRG5 mice (light grey), and Lck-Cre+; Mad2f/f; p53f/f mice (brown). Copy number changes of PRG5 mice were determined using whole-genome sequencing. Averaged copy number from single-cell whole genome sequencing (as shown in Figure 1d) is shown for tumors from Mad2 mice. Genomic Identification of Significant Targets in Cancer (GISTIC) 2.0 analysis showing significant aneuploidies across all tumors is shown at the bottom. The similarity of individual tumors to the GISTIC 2.0 output is presented in the green heatmap (see Supplementary Table 4 for similarity values). Scaled tumor weights are presented (ND – not determined).

Multifocal tumor analyses showed little DNA variation among different tumor regions in spontaneous p53-/- thymic lymphomas of PRG5 mice; however, tumors driven by transient CIN showed clear clonal DNA differences, correlating well with RNA expression and TCR sequencing data (Supplementary Figure 10a-c). Multifocal biopsy RNA expression analyses revealed high correlation between tumors driven by transient CIN, in contrast to the more heterogeneous spontaneous p53-/- tumors (Supplementary Figure 10d).

Use of the Genomic Identification of Significant Targets in Cancer (GISTIC) algorithm to determine the most significant chromosomal events in advanced tumor groups (spontaneous, Plk4-transient CIN induced, and Mad2-chronic CIN induced thymic lymphomas) revealed chromosomes 4, 5, 12, 14, and 15 to be significantly increased (qValue=0.01). A similarity test (see Methods) of individual tumors to the GISTIC profile revealed that tumors forming following transient CIN had a higher similarity value to the GISTIC profile than the spontaneous p53-/- tumors (Figure 4 and Supplementary Table 4). Thus, transient CIN (and chronic CIN to a lesser extent) accelerated the formation of a specific aneuploidy profile to which spontaneous tumors only displayed partial similarity.

### Aneuploidy profile selection occurs early in lymphoma development driven by transient CIN

Multifocal biopsy analysis of thymic lymphomas suggested that aneuploidy selection occurs early during tumor development (as different regions in the tumor present almost identical chromosome alterations and similar to the GISTIC profile) (Supplementary Figure 10). To critically test this, we used scWGS to determine the temporal dynamics of chromosome contents. Sequencing of early tumors (<500mg, with mice lacking any clinical signs such as weight loss or dyspnea) or late tumors (Supplementary Figure 11 and Figure 5a) revealed that chromosome 4, 5, 14, and 15 gains in CIN-induced tumors (both early and late) were present in almost every cell. Consistent with early selection of an aneuploidy profile, this outcome contrasted with similar analyses of spontaneous tumors in p53-/- mice (Supplementary Figure 11). Early tumors forming following transient CIN showed a very high similarity to the GISTIC profile identified in late tumors, indicating that optimal tumorigenic clones form early under this condition, and in contrast to spontaneously formed tumors (p53-/- mice) and tumors forming under chronic CIN (Mad2-/- mice) (Figure 5a).

**Figure 5.**
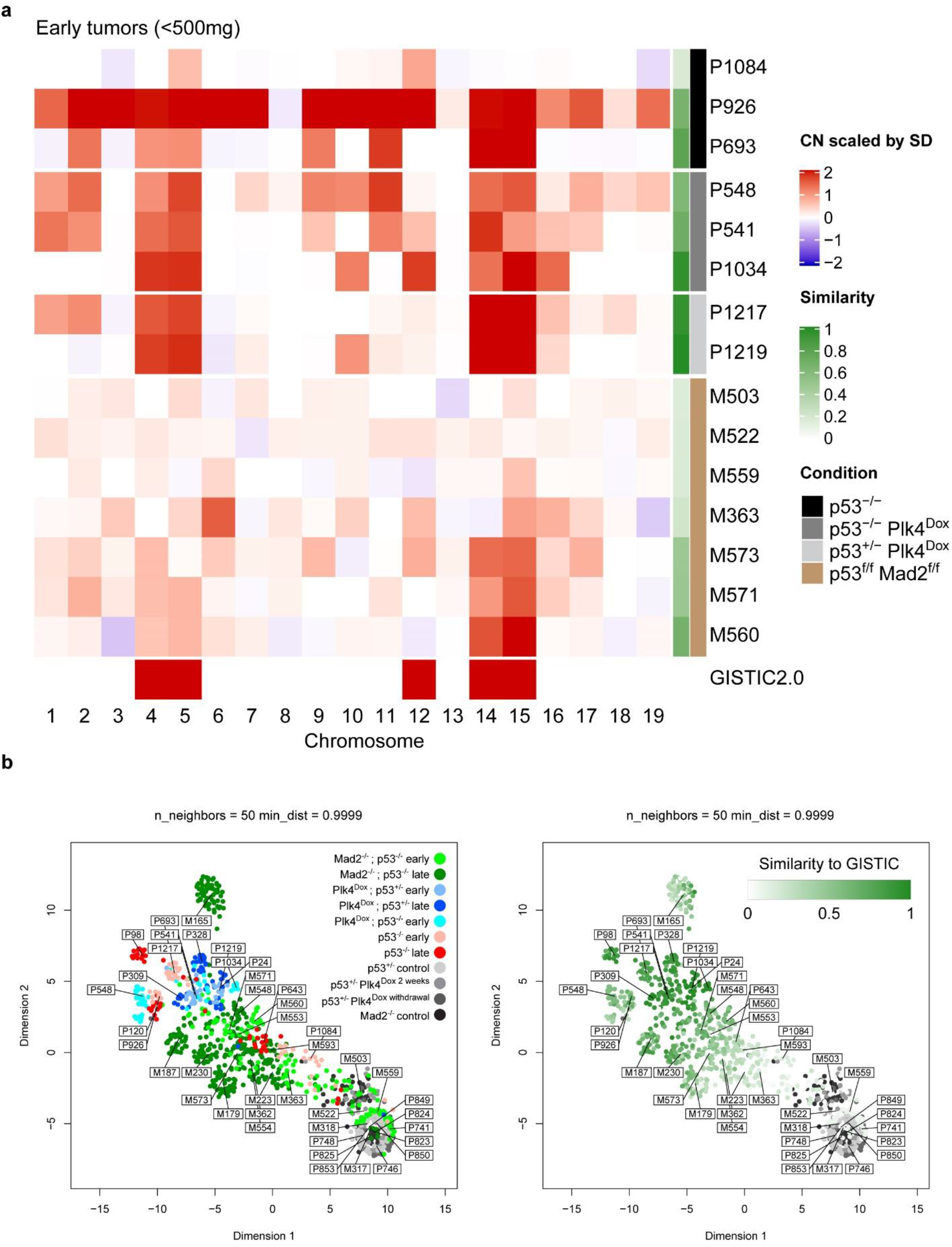
Transient CIN selects for a specific aneuploidy profile early during tumor development. (**a**) Heatmap showing averaged DNA copy number changes in early tumors (<500mg) from non-induced p53-/- PRG5 mice (black), two-weeks doxycycline treated (at the age of 30 days) p53-/- PRG5 mice (dark grey) and p53+/- PRG5 mice (light grey), and Lck-Cre+; Mad2f/f; p53f/f mice (brown) as determined using single-cell whole genome sequencing. Single cell data of tumors from PRG5 mice is presented in Supplementary Figure 11, and of Mad2 mice in Figure 1c. Genomic Identification of Significant Targets in Cancer (GISTIC) 2.0 analysis showing significant aneuploidies across all tumors is shown at the bottom. The similarity of individual tumors to the GISTIC 2.0 output is presented in the green heatmap. (**b**) left: Uniform Manifold Approximation and Projection (UMAP) analysis of DNA copy number changes in control (grey shades), p53-/- non induced PRG5 tumors (red shades), doxycycline induced PRG5 tumors (blue shades), and Mad2 tumors (green shades). Right: Embedding of the GISTIC 2.0 similarity score for each sample shown in the UMAP plot on the left. Control samples were not used for the GISTIC 2.0 analysis and appear in grey shades.

The most striking difference between genomes of tumors developed under chronic CIN or following transient CIN was the selection of chromosomes 4, 5, 14, and 15 gains apparently accumulating in single cells during early tumor development. Transient CIN-driven and spontaneous tumors showed significantly higher copy-number score (and higher aneuploidy) but lower copy-number score variability than chronic CIN driven tumors (Supplementary Figure 12). Using the Uniform Manifold Approximation and Projection (UMAP) algorithm, we determined the spatial distribution of high-dimensional data obtained from scWGS (Figure 5b). UMAP separated cell populations according to their tumor of origin (sample cell types were not given as an input - Figure 5b). A transition from euploid non-cancer cells to terminal tumors was observed, with early Mad2-/- tumors as well as p53-/- tumors containing cells within this spectrum. Plk4 driven tumors were localized with some p53-/- and terminal Mad2-/- tumors having spatial proximity. When marking the single cells with their respective GISTIC similarity score, UMAP separated single cells according to their similarity with a visible gradient from euploid cells to a cluster of cells with high similarity where both early and terminal Plk4 tumors localized. Thus, a very specific selection occurs in early tumors following transient CIN, a process which chronic CIN counteracts.

### Aneuploidy selection generates a gene expression profile of human cancers

Gene expression analysis performed in correlation with the similarity index of each tumor in PRG5 mice (see Methods for detailed analysis pipeline plus Supplementary Figures 13 and Supplementary Table 5) revealed significant enrichments of genes related to cell cycle progression and to pathways allowing cells to overcome oncogene-induced senescence and replication stress. This outcome raised the possibility that aneuploidy selection might provide higher tolerance to oncogene stress (due to overexpression of Myc, for example).

We examined the relevance of this proposed gene expression profile to what has been found in human cancers in the TCGA database (Supplementary Figure 14). This revealed that 11/24 of the cancers in this cohort substantially acquired a gene profile similar to that in tumors from transient CIN (Supplementary Figure 14a). This gene profile was highly correlated with increased MYC expression in 13/34 of the TCGA cancers (Supplementary Figure 14b) and was mutually exclusive between the two groups (with the exception of four cancer types: LUAD, LUSC, STAD, and UCEC), indicating that some tumors develop this profile in general, and some only after Myc is overexpressed.

## Discussion

In his prophetic monograph, Boveri laid the foundations for cancer research years to come^29^. In the current study, we tested Boveri’s hypothesis suggesting that an abnormal chromosome content can lead to cancer. We find that a very short timeframe of CIN is sufficient to trigger tumor initiation and acceleration in p53 heterozygous and null mice, respectively. The fact that tumor latency in p53 heterozygous mice experiencing transient CIN at an early age is almost identical to spontaneous tumor formation in p53 null mice suggests that the events responsible for thymic lymphoma in p53 null mice occur at the same time frame in which we induced CIN. Nevertheless, we find it surprising that tumor burden was higher in CIN-induced p53 heterozygous mice, as loss of chromosome 11 containing the wild type p53 gene is a prerequisite for thymic lymphoma formation in such mice. Considering that not all cells in the thymus lost the wild type p53 allele, this suggests that the rate of transformation of newly made p53 null cells is much higher in the CIN induced p53 heterozygous mouse, leading to multiple tumors forming.

We also found that p53 null mice experiencing transient CIN have accelerated onset of thymic lymphomas, at least partly due to increased clonal burden. Importantly, it was reported that thymic lymphomas in p53 null mice can form from more than one tumor initiating cell^30^, and our study shows that transient as well as chronic CIN drives this further. Tumors from both p53 heterozygous and null mice experiencing CIN at an early age show similar genomic landscape in which chromosomes 4, 5, 14, and 15 are gained. Strikingly, this genomic profile is in agreement with previous reports of thymic lymphomas in mice with Lck-Cre+ Mps1 mutation^10^, and also with thymic lymphomas derived from Mad2 knock out in this study and an earlier report^26^, although these works generate CIN through completely different mechanisms, and do so chronically.

In our multifocal biopsy DNA analyses, we found supporting evidence that the genomic landscape identified is found in a single cell, originating very early during tumor formation, as different regions in the tumor contain almost identical genome landscapes. Our data provides evidence for early and fast catastrophic aneuploid events already occurring during the two-week induction of CIN, consistent with the idea of the ‘Big Bang’ model proposed for human breast and colon cancers^31,32^. We propose that it is in this timeframe that multiple optimal and sub-optimal tumorigenic cells form, and that the optimal clones contain the aforementioned genomic landscape which is detected also in the developed tumor. Intriguingly, we observe the same catastrophic aneuploid events emerging in developing Lck-Cre; Mad2; p53 tumors, however they appear to struggle reaching the optimal karyotype, probably due to the constant CIN pressure, highlighting transient CIN as a more efficient driver of tumorigenesis.

Our data are also in agreement with data showing that aneuploidy is not well tolerated in normal cells^13,14^ as in thymic tissue it disappears after removing CIN. In p53 null mice spontaneously developing thymic lymphomas, a similar genomic landscape appears to be forming through an extended trial and error of ongoing clonal evolution, and probably given time, or examination of enough tumors (as the case of the multifocal biopsy from one of these mice, see Supplementary Figure 10), such a genomic landscape would appear. We present evidence for tissue specific aneuploidy profile that optimizes malignant transformation and propose that such optimal genomic landscapes exist for each tissue and cell type, with evidence for this already seen through our knowledge of recurrent chromosome abnormalities and of the involvement of chromosome instability^26,33,34^. Importantly, thymic lymphomas forming in developmentally arrested RAG2 deficient cells present a completely different genomic landscape lacking gains of chromosomes 1/4/5/14/15 but containing a unique rearrangement of chromosome 9qA4-5.3^35^, suggesting that even within the same tissue context, cancer genome evolution is also dependent on the genetic background of the cell. Our work shows that clonal evolution is operative in the development of thymic lymphoma and is accelerated under transient CIN.

## Acknowledgments

This work was funded by a grant from the US National Institutes of Health (R35 GM122476 to D.W.C.). D.W.C. receives salary support from the Ludwig Institute for Cancer Research. We thank Andrew Shiau for providing access to the CQ1 spinning disk confocal system. We thank Dr. Nissi Varki (University of California at San Diego Pathology Core) for histology sample analysis. We thank Kristen Jepsen from the UC San Diego IGM Genomics Center for help with whole genome DNA sequencing (National Institutes of Health SIG grant #S10 OD026929). We thank Stefan Kessler with help uploading the raw sequencing data to public databases.

## Author contributions

O.S. and D.W.C. conceived the project and wrote the manuscript. O.S. designed, performed, and analyzed the experiments. B.B. performed and analyzed the experiments related to samples from the Mad2 mouse cohort. Y.W. assisted in maintaining the Plk4 mouse cohort and helped with breeding and samples collections. D.H.K. assisted with live cell imaging. M.M. helped with tumor sample processing and imaging. M.A.D. helped with sample processing for RNA expression analysis. J.A. helped with maintenance of Plk4 mouse colony, and with RNA library preparation for RNA sequencing. O.Z. helped with analysis of Plk4 cohort RNA sequencing analysis. A.M., R.W., R.S., and K.F. helped analyzing RNA sequencing, whole-genome DNA sequencing, and single cell whole genome DNA sequencing analysis. D.C.J.S. helped preparing samples for single cell whole genome DNA sequencing. B.V. helped generating the Plk4 mouse cohort. All authors provided input on the manuscript and D.W.C., F.F., and O.S. supervised all aspects of the work.

## Methods

### Cell culture

Mouse embryonic fibroblasts (MEFs) were derived as previously described^36^. Briefly, E13.5 embryos were washed with PBS and head, liver, and tail were removed. Embryos were minced in 0.05% trypsin (Gibco) and incubated in 37°C for 15 minutes. Dissociated cells were plated in DMEM media (Gibco) supplemented with 15% fetal bovine serum (Omega Scientific), 50μg/ml penicillin and streptomycin (Gibco), 2mM L-glutamine (Gibco), 1μM 2-mercaptoethanol (Sigma), 1mM sodium pyruvate (Gibco), and 0.1mM non-essential amino acids (Gibco). Cells were maintained at 37°C, 5% CO2, and 3% O2. Doxycycline (Sigma) was dissolved in water and used at a final concentration of 1μg/ml. Thymic lymphoma cells were derived as previously described^37^. Briefly, dissected tumors were dissociated using bent needles, and cells were plated in RPMI 1640 with 25 mM HEPES, 200 mM L-glutamine (Lonza), and supplemented with 10% fetal bovine serum (Hyclone), 1% penicillin and streptomycin (Gibco), 1% non-essential amino acids (Gibco), and 55mM 2-mercaptoethanol (Sigma).

### Mice

Generation of doxycycline inducible Plk4 mouse with the tetracycline responsive Plk4-YFP gene inserted downstream of the Col1a1 locus and the M2-rtTA gene inserted into the ROSA locus was previously described^23^. To generate the PRG5 mice, Plk4 inducible mice were crossed with centrin-GFP mice^27^ and with mice carrying a knockout allele of p53^28^. Mice homozygous for the Plk4 transgene and for the M2-rtTA gene and heterozygous for p53 were crossed with mice homozygous for centrin-GFP and heterozygous for p53 Mad2 to generate cohort mice heterozygous for Plk4, M2-rtTA, and centrin-GFP, and homozygous or heterozygous for p53. Doxycycline was administered for indicated times through mouse diet containing 0.625 g/Kg Doxycycline Hyclate (Envigo, TD.08541). Genotyping performed on tail DNA was done using the following primers: Plk4-YFP - CACAGGAACAGGCGTCTCTTCAAGTC and GTGCAGATGAACTTCAGGGTCAGCTTG; rtTA - AGGAGCATCAAGTAGCAAAAGAG and AAGAGCGTCAGCAGGCAGCA; centrin-GFP - GACAAGCAGAAGAACGGCATCAAGGTG and CTTGCTTCTGATCCTCAGTGAGCTC; p53 - ACAGCGTGGTGGTACCTTAT and TATACTCAGAGCCGGCCT and TCCTCGTGCTTTACGGTATC; Rosa – AAAGTCGCTGAGTTGTTAT and GCGAAGAGTTTGTCCTCAACC and GGAGCGGGAGAAATGGATATG; Col1a1 – CCAGCTTCACCAGTTCAATCATCC and CAGTCCCTGTTTCTGCTGCTTGAATC. Mice were housed and cared for in an Association for the Assessment and Accreditation of Laboratory Animal Care-accredited facility, and all animal experiments were conducted in accordance with Institutional Animal Care and Use Committee-approved protocols. Lck-Cre; Mad2f/f; p53f/f mice were previously described^26^. Genotyping of Lck-Cre; Mad2f/f; p53f/f mice was performed as described previously^26^. Mice were housed and experiments were conducted according to Dutch law and approved by the Central Committee Animal experiments (CCD, permit AVD105002016465). Tissue sections from formalin fixed paraffin embedded (FFPE) tissues and from cryopreserved tissues were collected as previously described^22^.

### Centrosome enumeration

Centrosomes visualized by the centrin-GFP marker in PRG5 derived mouse embryonic fibroblasts grown in chamber slides (ibidi) were imaged (1μM x 10, X40/1.35NA) using the DeltaVision elite system (Applied Precision). Centrosomes visualized by the centrin-GFP marker in in tissue cryosections were imaged (at 0.2μm Z-sections with a Nikon 100× APO TIRF 1.49 NA objective) using the Nikon A1 scanning confocal microscope operated with NIS-Elements (Nikon).

### Live cell imaging

PRG5 derived mouse embryonic fibroblasts were seeded in 96-well in CELLSTAR μClear 96-well plate (Greiner bio-one). Cells were stained with DNA SiR (Spirochrome) and three hours later were imaged using a CQ1 spinning disk confocal systems (Yokogawa Electric Corporation) with a x40 magnification at 37°C and 5% CO2. To induce Plk4 overexpression, doxycycline (Sigma, 1μg/ml) was added 1 hour prior to imaging. Live imaging of 8 × 3-μm z-sections was conducted for 24 hours. Image acquisition and data analysis were performed using CQ1 software and ImageJ, respectively. For time lapse imaging of Lck-Cre Mad2f/f p53 T-ALL cells, primary T-ALL derived cell lines were transduced with H2B-Cherry using retroviral transduction as described previously^38^, cultured in LabTek imaging chambers (Nunc) and imaged on a DeltaVision Elite microscope (Applied Precision). Mitotic abnormalities were quantified by manual inspection of the movies.

### Quantitative real-time PCR

RNA from PRG5 derived mouse embryonic fibroblasts was extracted using the Nucleospin RNA kit (Macherey Nagel). Mouse tissues were homogenized using a mechanical tissue homogenizer in Trizol reagent (Invitrogen) and RNA was extracted according to the manufacturer guidelines. cDNA was prepared from 1 μg total RNA using the high-capacity reverse transcription kit (ABI) according to the manufactures’ instructions. Quantitative real-time PCR was done in triplicates, using iTaq Universal SYBR green (Bio-Rad) and a CFX384 real-time PCR machine. For detection of Plk4, the following primers were used – GGAGAGGATCGAGGACTTTAAGG and CCAGTGTGTATGGACTCAGCT. The following primers were used as house-keeping control genes – Rsp9 – GACCAGGAGCTAAAGTTGATTGGA and GCGTCAACAGCTCCCGGGC; Actg1 – TGGATCAGCAAGCAGGAGTATG and CCTGCTCAGTCCATCTAGAAGCA.

### Protein analysis

Total protein from PRG5 mouse tissues (10mg biopsies) were extracted in 500μL X2 Laemmli sample buffer and 10μL from each sample was loaded in 10% acrylamide gel for SDS/PAGE separation. Proteins were transferred to a nitrocellulose membrane, blocked with 5% mils in tris-buffered saline and 0.1% Tween-20 (TBS-T), and incubated overnight with the following primary antibodies: anti-GAPDH (cell signaling, #2118, 1:10,000), Myc (Abcam, ab32072, 1:1,000). Immunoblots were washed with TBS-T and incubated with HRP-conjugated secondary antibodies (GE Healthcare, 1:5,000) for one hour in room temperature before development in films. For quantification of mitotic cells, cultured T-ALL cells were exposed to nocodazole (Sigma) or DMSO (Sigma) for 6 hours fixated in 70% ethanol, washed with PBS and blocked in blocking buffer (0.05% Tween (Sigma)/ 2% BSA (Sigma) in PBS) and stained with FITC-conjugated MPM2 antibody (Upstate) for 2 hours. Cells were washed in PBS and resuspended in staining buffer to label DNA was stained using. FACS staining buffer (20 ug/ml propidium iodide (Sigma), 0.2 mg/ml RNAseA (Sigma) in PBS). Cells were analyzed on a FACSCanto analyzer (BD) and quantified using FlowJo software (BD).

### Single cell whole genome DNA sequencing

Single cells from mouse thymuses and thymic tumors were isolated using flow cytometry sorting and prepared for sequencing as previously described^26^. Sequencing was performed using a NextSeq 500 machine (Illumina; up to 51, 77 or 84 cycles; single end). The generated data were subsequently demultiplexed using sample-specific barcodes and changed into fastq files using bcl2fastq (Illumina; version 1.8.4). Reads were afterwards aligned to the mouse reference genome (GRCm38/mm10) using Bowtie2 (version 2.2.4)^39^. Duplicate reads were marked with BamUtil (version 1.0.3)^40^. The aligned read data (bam files) were analyzed with AneuFinder (Version 1.14.0)^41^. Following GC correction and blacklisting of artefact-prone regions (extreme low or high coverage in control samples), libraries were analyzed using the dnacopy and edivisive copy number calling algorithms with variable width bins (binsize: 1 Mb; stepsize: 500 kb) and breakpoint refinement (R= 20, confint = 0.95; other settings as default). Results were afterwards curated by requiring a minimum concordance of 95% between the results of the two algorithms. Libraries with less than five reads per bin per chromosome copy (~ 25,000 reads for a diploid genome) were discarded. AneuFinder gave unexpected results for sample P309. About half of the libraries showed an average copy number of 1.5 and the other half an average copy number of 3 (two times as high). Examination of the model results showed poor fits for the first group of libraries. Sample P309 was therefore reanalysed with the developer version of AneuFinder (Version 1.7.4; from GitHub) using a minimum ground ploidy of 2.5 (min.groun.plody=2.5) and a maximum ground ploidy of 3.5 (max.ground.ploidy=3.5). Results were subsequently curated as described above. As a final step, breakpoints that were located within 5 Mb from each other (across libraries) were grouped and centered using custom made R functions (R version 3.6.3, https://www.R-project.org/) to prevent the heterogeneity scores to reach unlikely high values just because breakpoints are off by a few bins.

### Genome-wide karyotype measures

Copy number change score (CN change score): For each cell this is calculated as the average absolute difference between the observed copy number of each bin and the expected copy number of each bin (euploid genome). Bins have variable width and therefore a weighted average was used. The score of the sample is calculated as the average score of all cells. Heterogeneity score: For each bin this is calculated as the proportion of all pairwise comparisons (Cell 1 vs. Cell 2, Cell 1 vs. Cell 3, etc.; Total number = n*(n-1) / 2) that show differences in copy number. The score of the sample is calculated as the average score of all bins. Mean number of transitions per Mb (Structural score): For each cell this has been calculated as the total number of transitions (or breakpoints) divided by the total genome length (sum of bin widths). The score of the sample is calculated as the average score of all cells.

### Uniform Manifold Approximation and Projection (UMAP)

The UMAP function of the R package umap (version 0.2.6.0; from published preprint: Leland McInnes, John Healy, James Melville (2020). UMAP: Uniform Manifold Approximation and Projection for Dimension Reduction. ArXiv 1802.03426.) was used to compute the manifold approximation and projection of all single cell DNA sequencing samples described in this paper (Figure 1b - d, Figure 2h, j, l and Supplementary Figure 11; Arguments: random_state=1, n_neighbors=50, min_dist=0.9999). A matrix containing the copy number calls of the 1 Mb bins of the autosomes of all samples was used as input. Chromosome X was excluded due to differences in gender.

### DNA fluorescent in-situ hybridization (FISH)

Dissociated splenic and thymic cells were fixed using methanol/acetic acid (3:1), washed with fixative three times, and kept in fixative at −20 °C until use. To make custom Trp53 BAC probes, the following BACs were ordered from bacpac (https://bacpacresources.org/): RP24-285L20 and RP23-243M15. BACs were isolated from 50ml bacterial cultures using the BACMAX Bac extraction kit (Epicentre). Isolated BACs were sonicated (x10 cycles of 15 seconds ‘on’ 45 seconds ‘off’ at constant intensity with power set to ‘3’, Branson sonifier 450) and labeled with TM-rhodamine Label-IT (Mirus). Labeled BAC probes were suspended in commercial chromosome paint probe for chromosome 11 (Metasystems). DNA-FISH was performed by applying probes onto samples and covering with a glass coverslip. Genomic DNA and probes were co-denatured at 75 °C for 2 minutes by placing slide on pre-heated metal plate. Samples were hybridized overnight at 37 °C in a dark humidified chamber. Slides were subsequently washed with 0.4× SSC at 72 °C for 2 min and rinsed in 2× SSC, 0.05% Tween-20 at room temperature for 30 s. Slides were then rinsed in PBS, counterstained with DAPI, and mounted using pro-long gold (Invitrogen). FISH images were acquired on a DeltaVision elite system (Applied Precision) at ×40 magnification (10 × 0.5 μm z-sections). Maximum intensity projections were generated using the softWoRx program.

### Preparation of DNA/RNA from tissue biopsies for sequencing

Tissues were snap frozen in liquid nitrogen and kept in −80 °C. Biopsies were taken from frozen tissues placed on a −80 °C chilled metallic stage. Stage and tools were cleaned between different tissues and biopsies to minimize cross contamination of DNA and RNA. Biopsies were homogenized using QIAshresdder (Qiagen) and DNA/RNA were prepared using the AllPrep DNA/RNA Mini Kit (Qiagen) according the manufacturer guidelines.

### TCR sequencing

TCR libraries were prepared using immunoSEQ kit mmTCRB according to the user manual (Adaptive biotechnologies). TCR libraries were sequenced using illumina Nextseq 500 (DNA Link facility), and TCR sequences were analyzed using the immunoSEQ Analyzer platform (Adaptive biotechnologies). Rare sequences (found at less than 1%) with 100% identity to dominant sequences were filtered and are marked in Supplementary table 3.

### Whole genome DNA sequencing (WGS)

Library Preparation: 450 nanograms of Genomic DNA from each sample was fragmented by Adaptive Focused Acoustics (E220 Focused Ultrasonicator, Covaris, Woburn, Massachusetts) to produce an average fragment size of 350 basepairs (bp). Fragmented DNA was purified using the Agencourt AMPure XP beads (Beckman Coulter, Fullerton, CA, USA) and sequencing libraries were generating using the KAPA Hyper Prep Kit (KAPA Biosystems, Wilmington, MA, USA) following manufacturer’s instructions using 4 cycles of amplification. The quality of the library was assessed using High Sensitivity D1000 kit on a 2200 TapeStation instrument (Agilent Technologies, Santa Clara, CA, USA). Sequencing was performed using the NovaSeq 6000 Sequencing System (Illumina, San Diego, CA, USA), generating 150 bp paired-end reads to obtain 10X coverage.

### RNA sequencing

cDNA libraries were prepared using the TruSeq Stranded mRNA Sample Preparation Kit (Illumina) according to the manufacturer guidelines. Then, cDNA libraries were sequenced on an Illumina HiSeq 4000 using single read, 50 cycle runs. FASTQ files were processed to assess quality by determining general sequencing bias, clonality and adapter sequence contamination. RNA sequencing reads were aligned to the mm10 mouse reference genome using STAR^42^. Gene expression levels TPM and raw counts were calculated by using RSEM^43^. The log2(TPM) values of selected TRBV transcripts were shown in a heatmap where the blue color stands for low expression while red for high expression. The heatmap is generated by ‘pheatmap’ package for the R program (R Core Team (2020). R: A language and environment for statistical computing. R Foundation for Statistical Computing, Vienna, Austria. URL https://www.R-project.org/).

### GISTIC 2.0 Similarity Index

CNVkit was used to determine DNA copy number from tumor WGS. We executed GISTIC 2.0 broad-level analysis using CNVkit segment files as input. Amplifications and deletions with q < 0.01 were kept, leaving chr 4,5,12,14,15 as significant amps. Segments were binned into 100,000 bp bins, each keeping their respective segment mean. Bin value z-scores were applied to the dataset, followed by an ecdf function. The resulting measure was then subtracted from 1 to create a “similarity index”, indicating a similarity to the “optimal clone” which is described by significantly amplified regions.

### CNV Heatmaps

R Bioconductor package ComplexHeatmap was used to plot the broad-level gistic results from the file “broad_values_by_arm.txt”.

### RNA-seq vs Normal Thymus

The R Bioconductor packages edgeR and limma were used to implement the limma-voom method for differential expression analysis. Lowly expressed genes were removed and then trimmed mean of M-values (TMM) normalization was applied. The experimental design was modeled upon experimental treatment of tumors (~0 + Treatment). The voom method was employed to model the mean-variance relationship in the log-cpm values, after which lmFit was used to fit per-gene linear models and empirical Bayes moderation was applied with the eBayes function.

### RNA-Seq using GISTIC 2.0 index as covariate

Samples’ similarity indices were used as continuous covariates in the RNAseq gene expression analysis using the limma-voom method to determine genes who’s expression significantly correlates with the similarity index. The experimental design was modeled upon the gistic index and batch (~gistic+batch).

### Data availability

RNA sequencing is deposited in the Gene Expression Omnibus (GEO) database, accession number GSE161728 (currently in private status). Whole-genome DNA sequencing is deposited in the Sequence Read Archive (SRA) database (currently in private status). Single-cell whole-genome sequencing is deposited in the European Nucleotide Archive (ENA) accession number PRJEB41176.

---

**Supplementary Figure 1.**
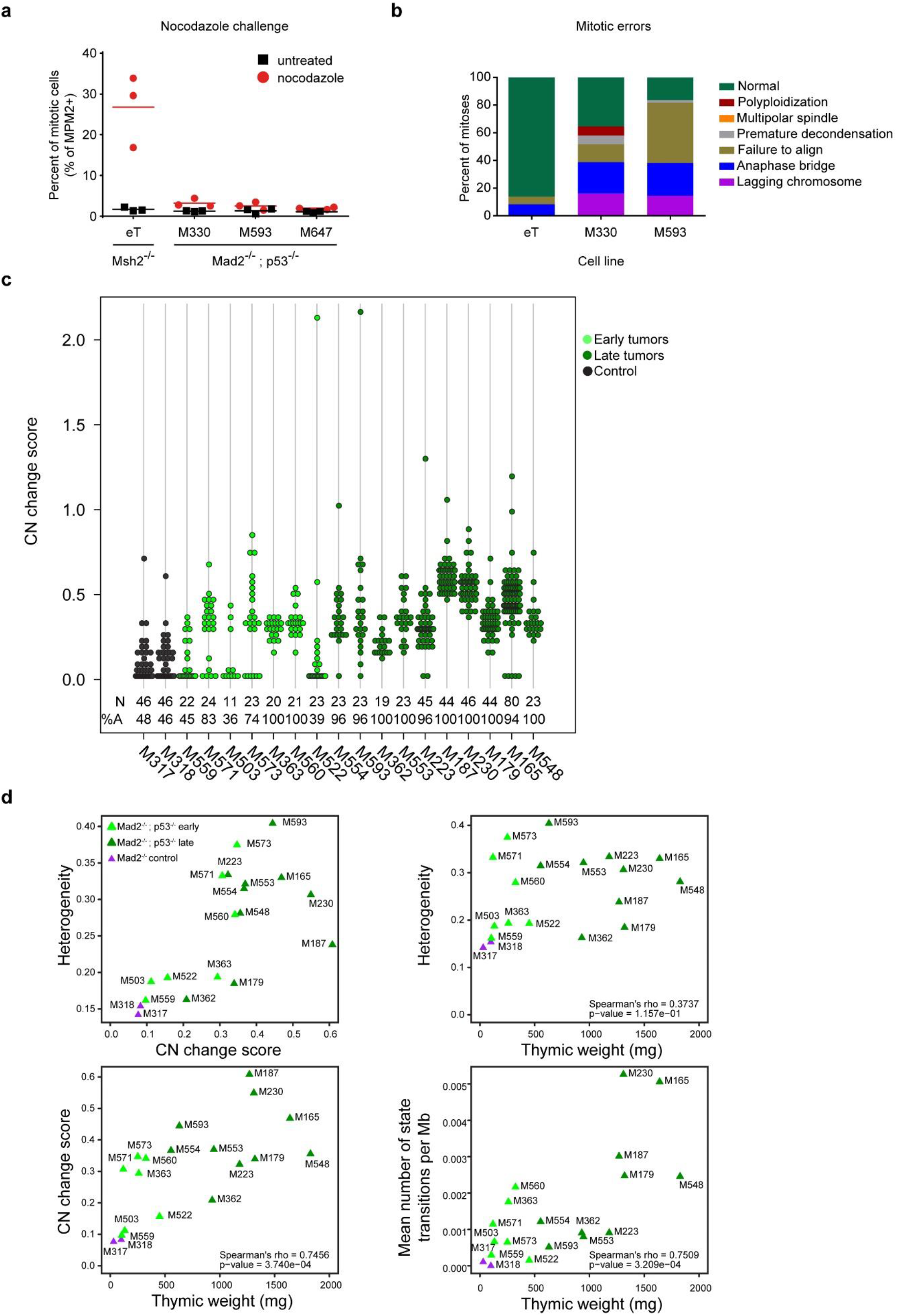
Mad2-deficient thymic lymphomas lack a functional SAC and display ongoing CIN. (**a**) Percentage of mitotic (MPM2+) cells in primary Mad2-/-; p53-/- or Msh2-/- T-ALL cultures with or without a nocodazole challenge for six hours. (**b**) Frequency of abnormal mitoses observed in Mad2f/f; p53f/f; Lcke primary T-ALL cultures as determined using live-cell time-lapse imaging after transduction with H2B-mCherry. (**c**) Analysis of copy number changes of the samples pf Figure 1b-d. The numbers below the dot plots indicate the total number of cells and the percentage of cells that has at least one whole chromosome gain or loss. (**d**) Genome-wide karyotype measures (heterogeneity score and CN [copy number] change score) for all samples shown in Figure 1b-d. Structural aberration scores (Mean number of state transitions per Mb) were defined by the number of copy number state transitions per Mb plotted against the weight of the thymus at time of harvest. Correlations were determined using Spearman’s rank-order test. See methods for details about the karyotype measures displayed.

**Supplementary Figure 2.**
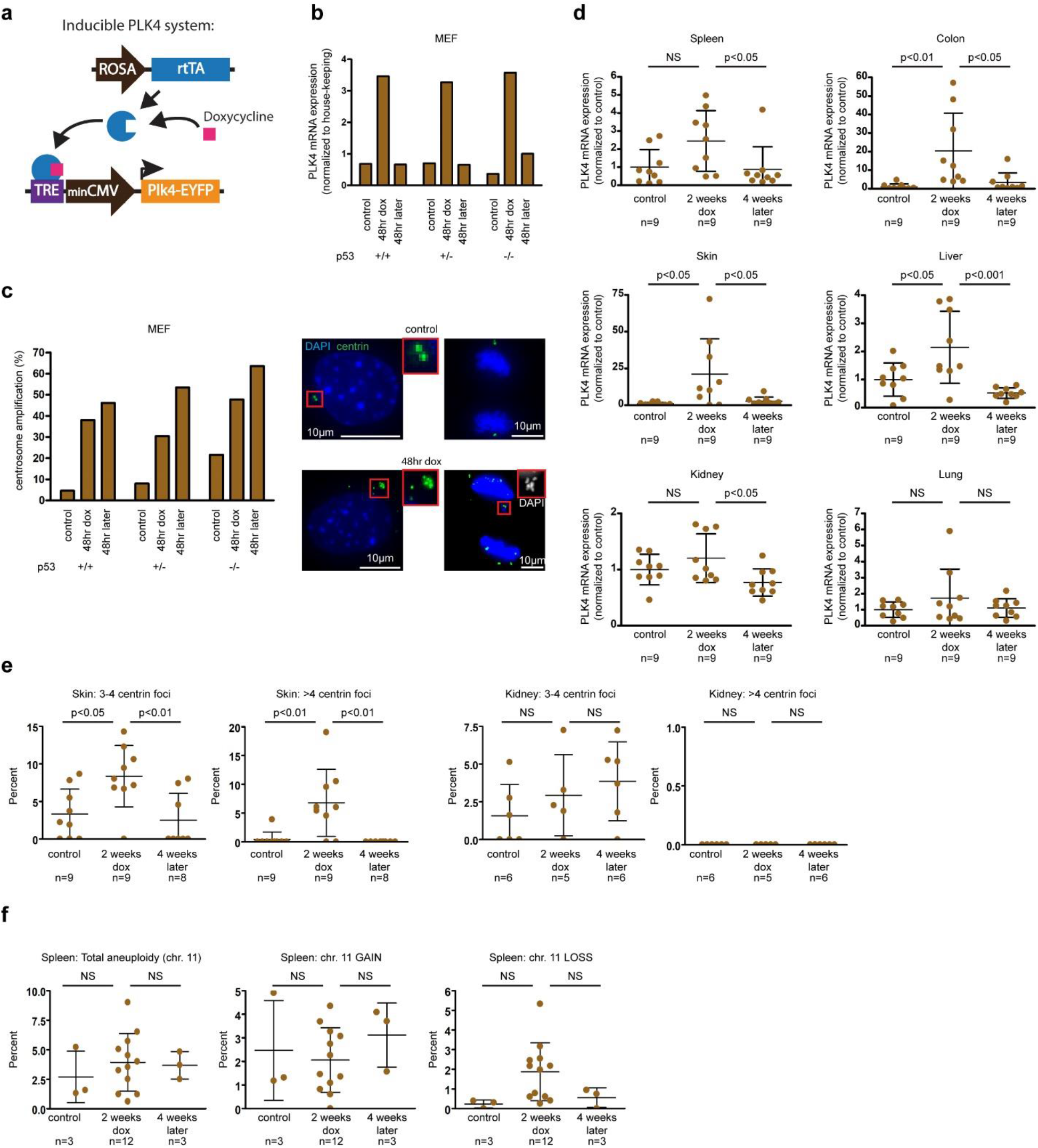
Effect of PLK4 overexpression in vitro and in vivo. (**a**) Inducible Plk4 expression using the tetracycline-on system. Doxycycline provided in mouse chow binds the reverse tetracycline-controlled trans-activator (rtTA, inserted in the Rosa locus) that together bind a Tet Response Element (TRE) upstream of Plk4 (inserted into the Col1A locus). (**b**) Plk4 mRNA levels in mouse embryonic fibroblasts derived from p53+/+, p53+/-, and p53-/- PRG5 mice, before (control) immediately after (48hr dox) and 48 hours after (48hr later) doxycycline administration. (**c**) Frequency of centrosome amplification as determined using centring-GFP in mouse embryonic fibroblasts derived from p53+/+, p53+/-, and p53-/- PRG5 mice, before (control) immediately after (48hr dox) and 48 hours after (48hr later) doxycycline administration. (**d-f**) Plk4 mRNA levels (**d**), measurement of centrin-GFP foci (**e**), and percent aneuploidy for chromosome 11 (using interphase DNA-FISH, **f**) in indicted tissues from PRG5 mice before (control) immediately after (2 weeks dox) and one month after (4 weeks later) doxycycline administration. Mean ± SD of indicated mice per group are presented. *p-values determined using one-way ANOVA with Tukey’s Multiple Comparison Test.

**Supplementary Figure 3.**
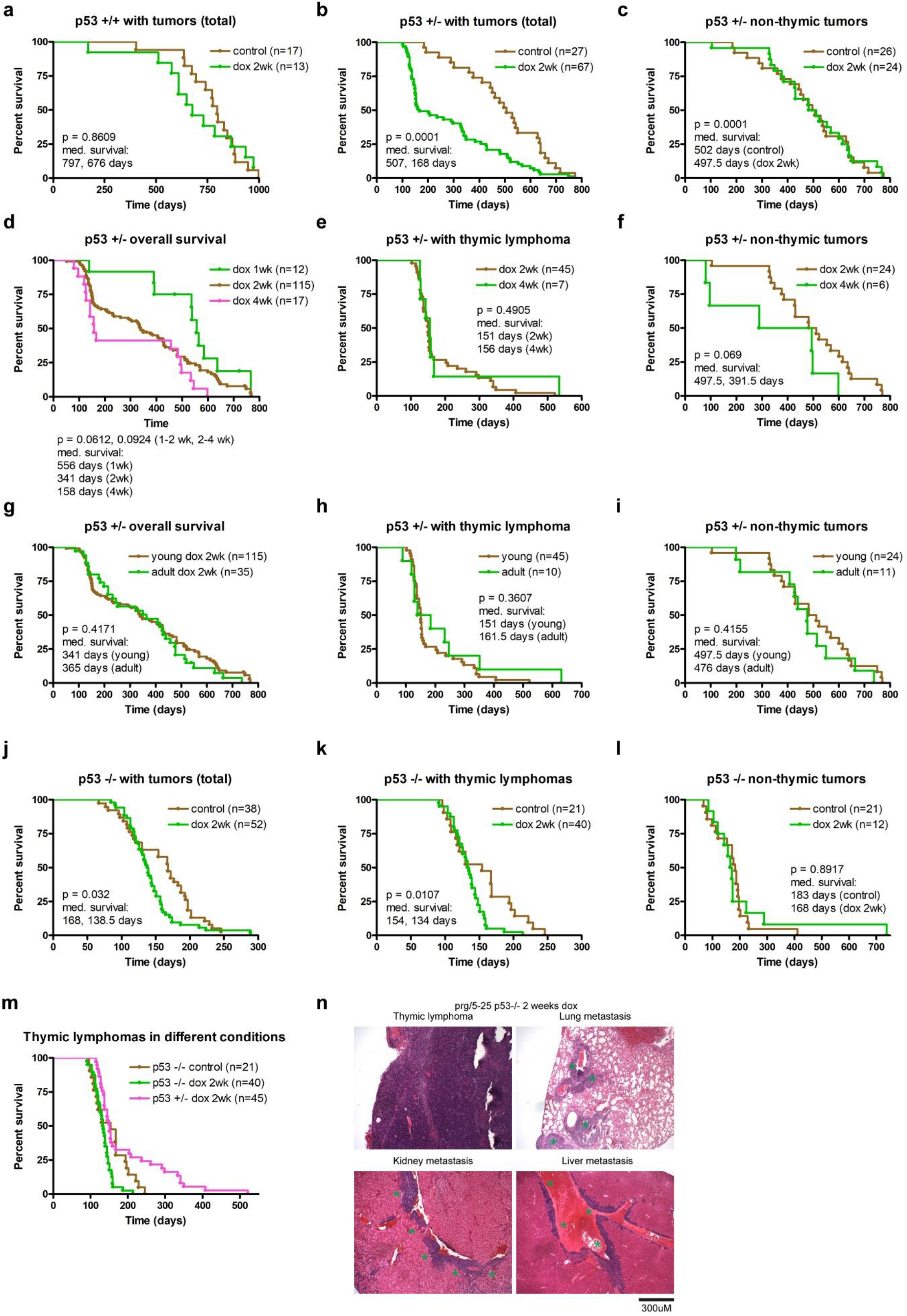
Effects of timing and duration of chromosome instability on tumor formation. (**a-m**) Survival (Kaplan-Meier) plots of PRG5 mice treated with the indicated doxycycline regimens (dox 1wk/2wk/4wk – treated with doxycycline for 1/2/4 weeks at the age of 30 days; young/adult dox 2wk – 2 weeks treatment of doxycycline starting at 30/100 days of age) and with the indicated p53 backgrounds (p53+/+, p53+/-, p53-/-). Overall survival and survival of mice bearing either thymic or non-thymic tumors are presented. Comparison of indicated number of mice done using log rank test. (**n**) Representative images of thymic lymphoma metastases from a 2-weeks Plk4 induced p53-/- PRG5 mouse. See Supplementary Table 1 for a histology analysis of tissues from a total of 104 mice.

**Supplementary Figure 4.**
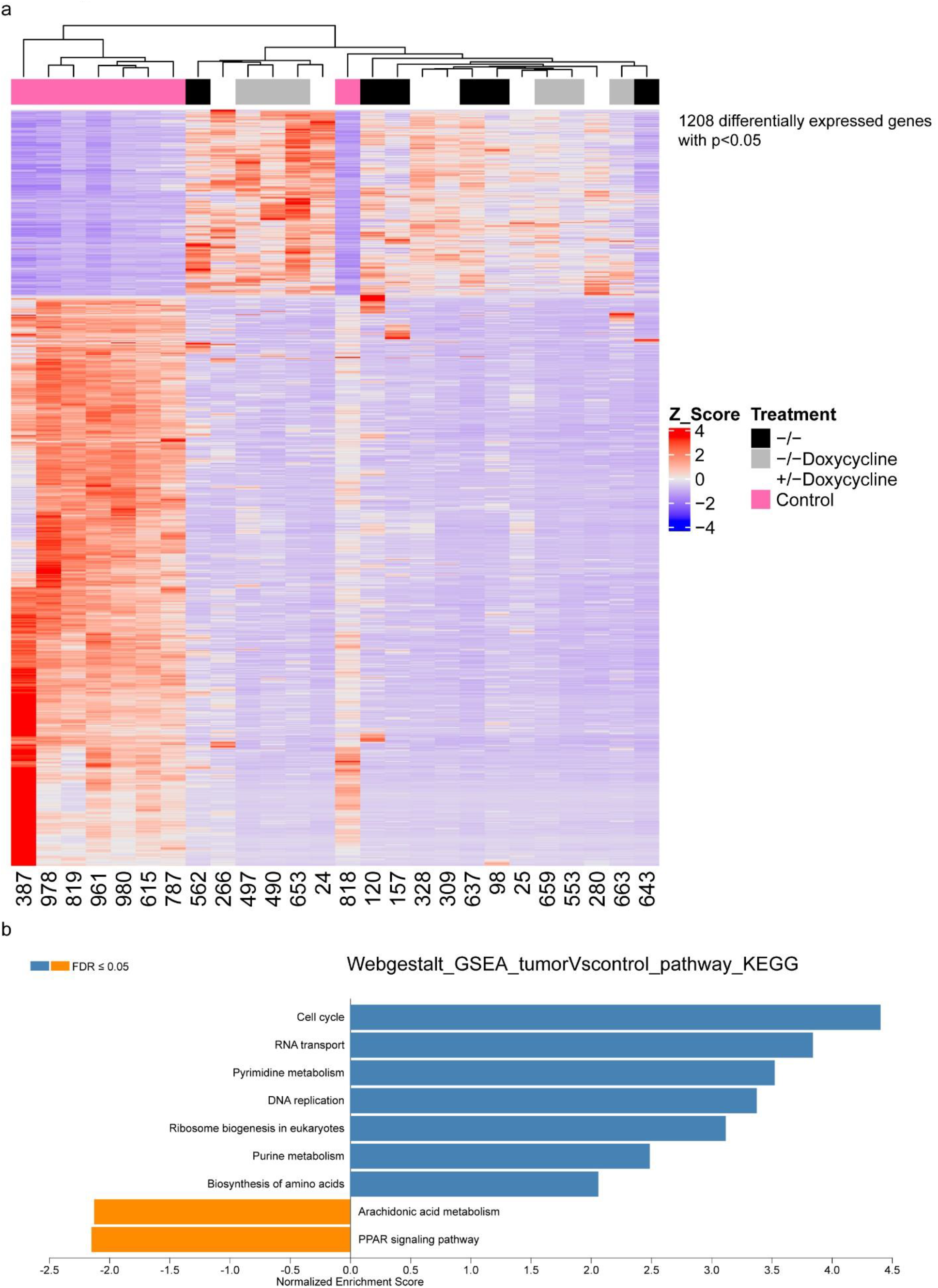
Differential gene expression analysis comparing thymic lymphomas and normal thymus. (**a**) Differentially expressed genes as determined using RNA sequencing between control normal thymuses and PRG5 derived thymic lymphomas. See Supplementary Table 2 for a complete list of the differentially expressed genes (**b**) Significantly enriched KEGG pathways in control thymuses and in PRG5 derived thymic lymphomas determined using Webgestalt GSEA analysis.

**Supplementary Figure 5.**
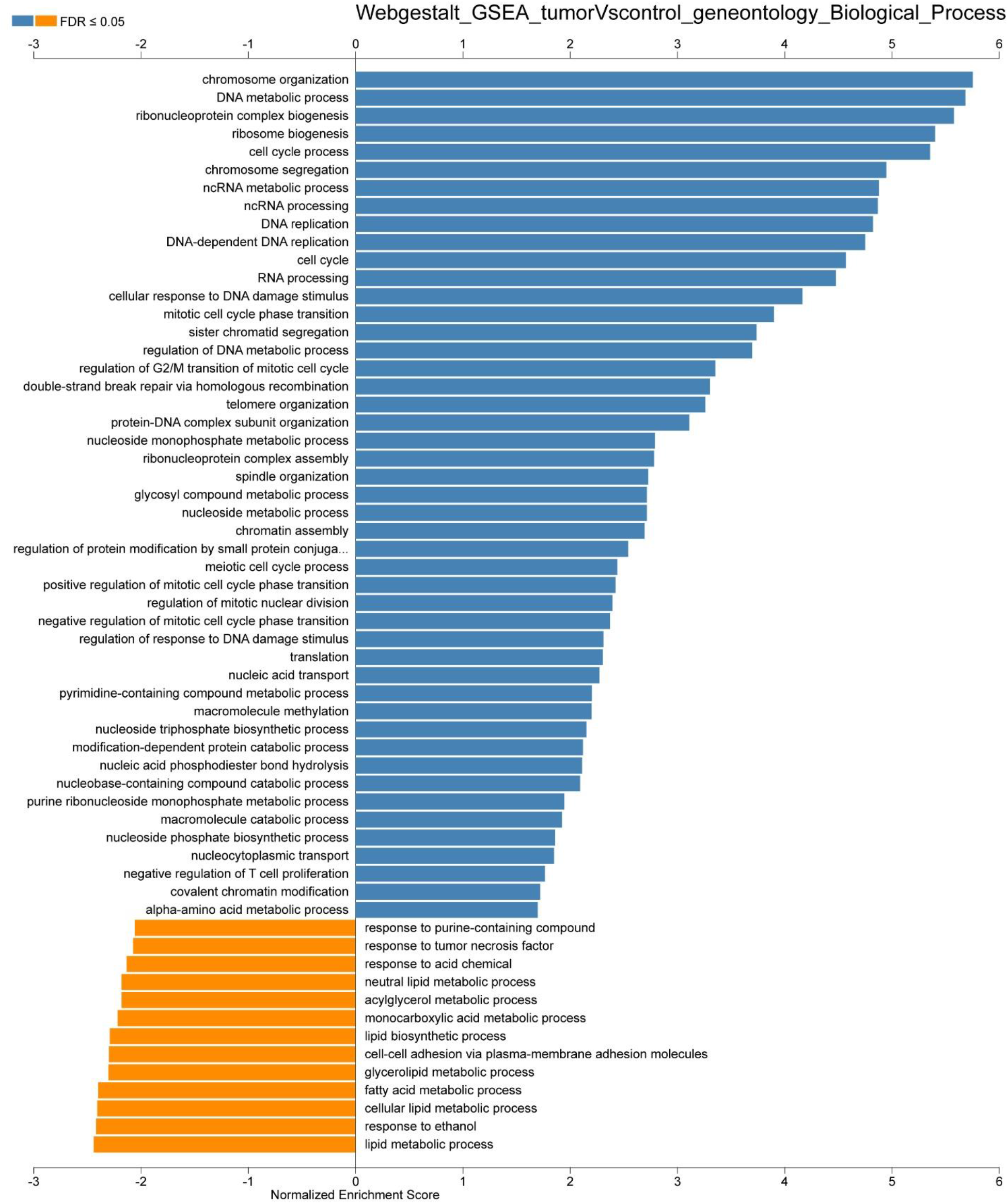
Pathway analysis reveals differences between thymic lymphomas and normal thymus. Significantly enriched Gene Ontology (Biological Processes) pathways in control thymuses and in PRG5 derived thymic lymphomas determined using Webgestalt GSEA analysis.

**Supplementary Figure 6.**
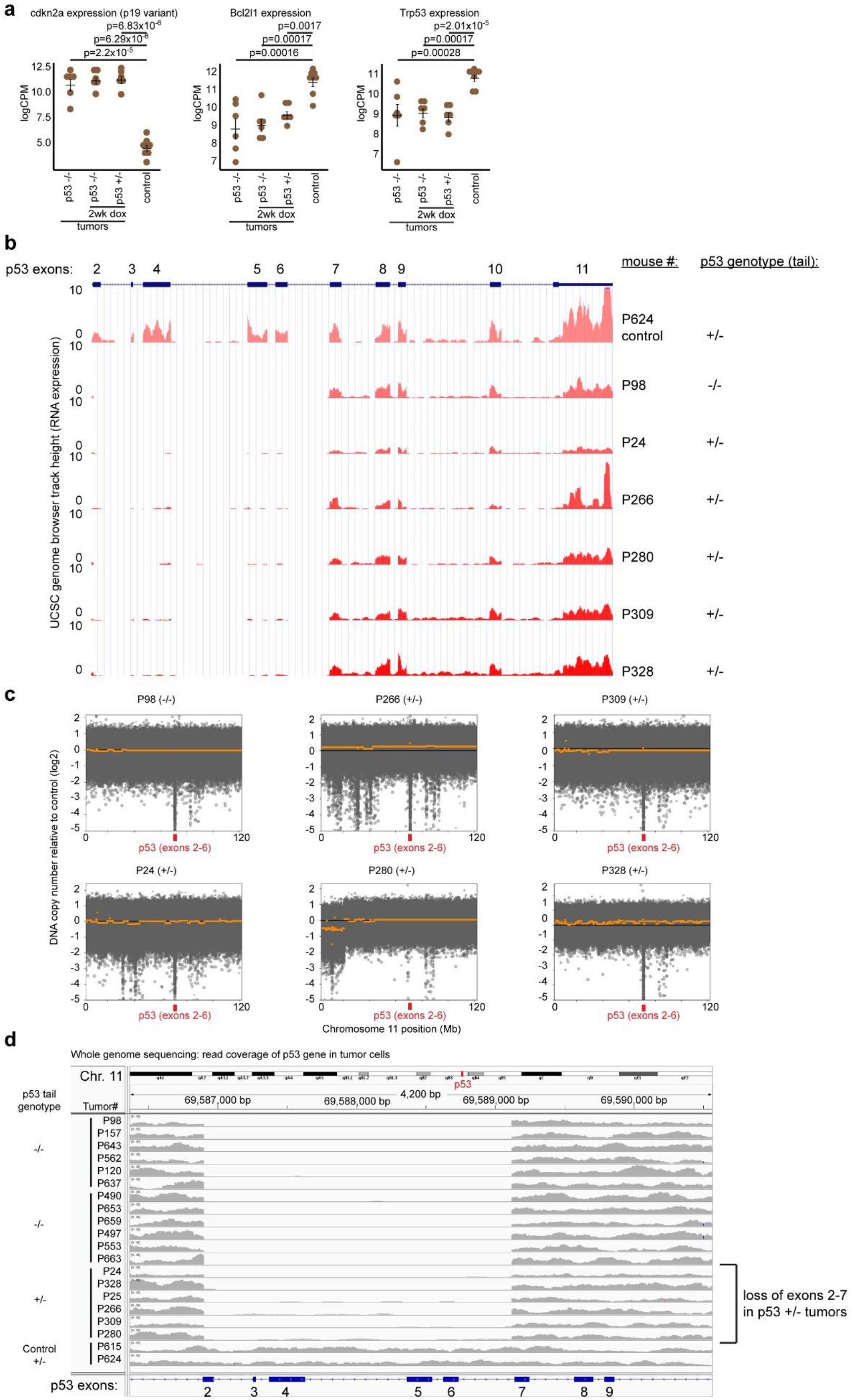
p53 LOH occurs through whole-chromosome missegregation without structural changes. (**a**) Comparison of mRNA expression levels (log[counts per million]) of indicated PRG5 tumors (n=6 tumors from each group) and control thymuses (n=8). Mean ± SD, and p-values calculated using one-way ANOVA are presented. (**b**) UCSC genome browser tracks showing RNA expression of exons 2-11 of p53 in indicated mice. (**c**) CNVkit derived DNA copy number plots of chromosome 11 in tumors from the indicated PRG5 mice. Black line represents the diploid control and the orange line represents the mean copy number of the indicated sample. The location of the *p53* gene is indicated in red. (**d**) Zoom on exons 2-6 of the p53 gene using Integrated Genome Viewer (IGV 2.3.97) showing number of reads in the indicated samples.

**Supplementary Figure 7.**
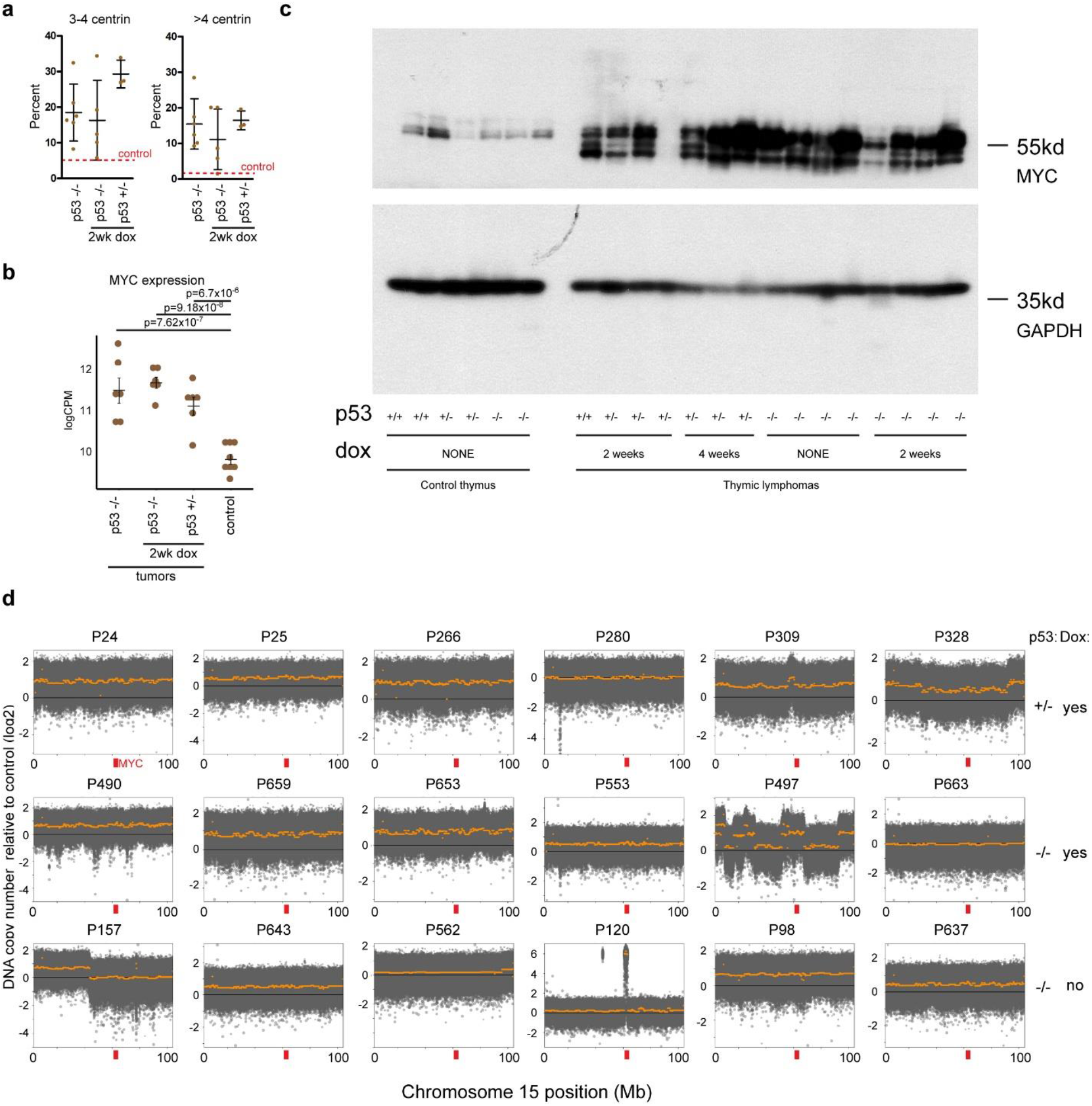
Common molecular events in spontaneous and induced lymphomas. (**a**) Measurement of centrin-GFP foci in thymic lymphomas from indicated PRG5 mice. Mean ± SD of n=6 (p53-/-), n=5 (p53-/- 2wk dox), and n=3 (p53+/- 2wk dox) are presented. Dashed red line represents the control shown in Figure 2f. (**b**) Comparison of mRNA expression levels (log[counts per million]) of indicated PRG5 tumors (n=6 tumors from each group) and control thymuses (n=8). Mean ± SD, and p-values calculated using one-way ANOVA are presented. (**c**) Myc protein levels as seen using SDS gel electrophoresis immunoblotting in the indicated control or tumor samples. GAPDH is shown as a loading control. (**d**) CNVkit derived DNA copy number plots of chromosome 15 in tumors from the indicated PRG5 mice. Black line represents the diploid control and the orange line represents the mean copy number of the indicated sample. The location of the MYC gene is indicated in red.

**Supplementary Figure 8.**
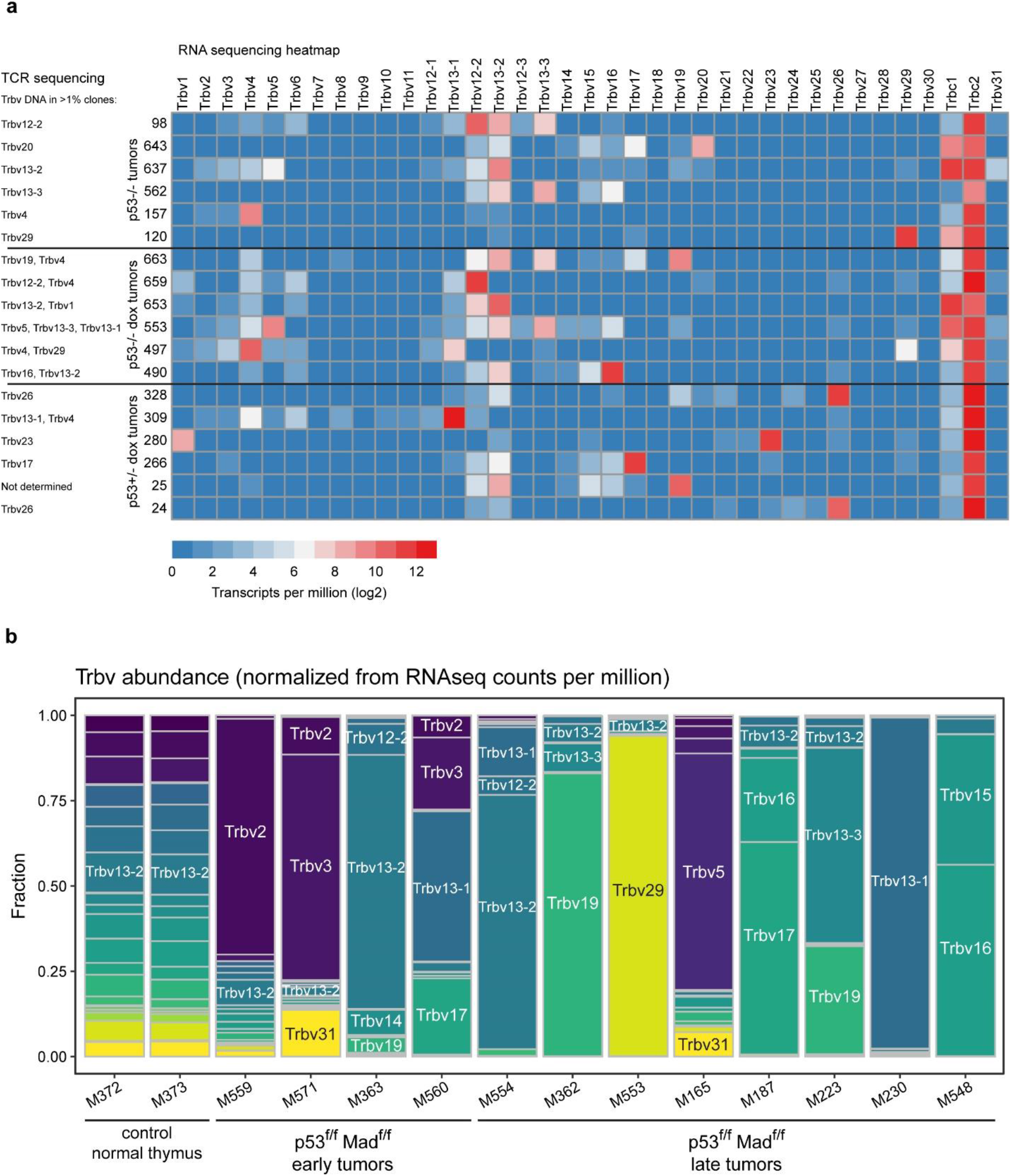
Tumor clonality derived from RNA sequencing data. (a-b) Expression of T cell receptor variable region genes (Trbv genes) as determined using RNA sequencing in tumors from (**a**) PRG5 mice and (**b**) Mad2 mice. Trbv genes with frequency > 1% in PRG5 tumors as detected using T cell receptor DNA sequencing (as shown in Figure 3j-k) is presented in **a**.

**Supplementary Figure 9.**
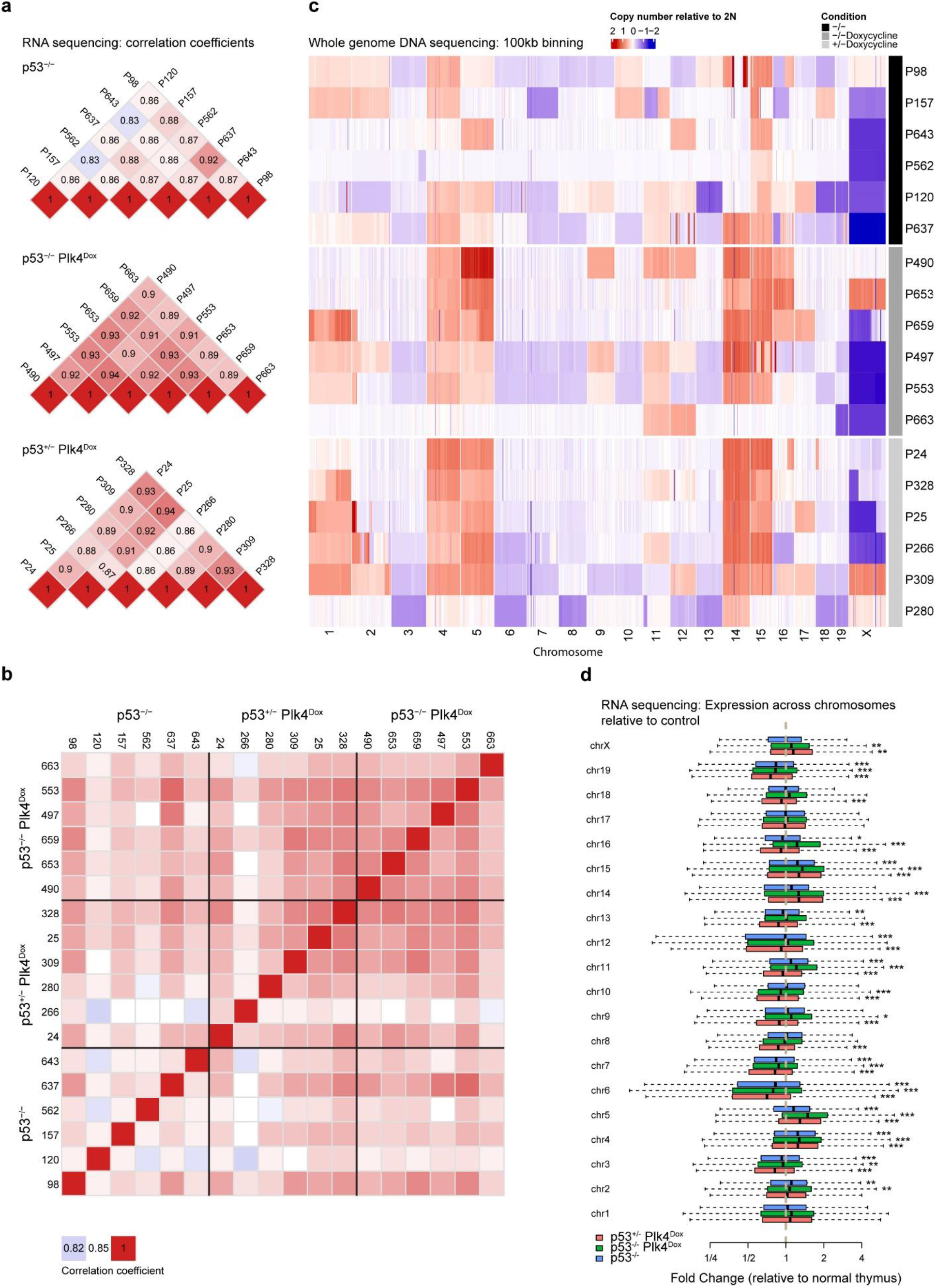
DNA and RNA levels in single biopsies of thymic lymphomas. (**a-b**) Heatmaps showing pair-wised Pearson correlation coefficients among samples on TPM values as determined using RNA sequencing of the indicated PRG5 tumor in (**a**) single biopsies or (**b**) multiregional biopsies. (**c**) Heatmap showing 100-kb DNA copy number changes in late tumors (>500mg) from non-induced p53-/- PRG5 mice (black), two-weeks doxycycline treated (at the age of 30 days) p53-/- PRG5 mice (dark grey) and p53+/- PRG5 mice (light grey) as determined using whole-genome sequencing. (**d**) The fold change of each gene between each tumor group and the control (n=6 for each group) was calculated, and boxplots of grouping all genes by chromosome for each comparison are presented. p-values (comparing expression levels from individual chromosomes between of each tumor group and the control) were determined using wilcoxon rank sum test, *: p<0.05; **:p<0.01 and ***:p<0.001.

**Supplementary Figure 10.**
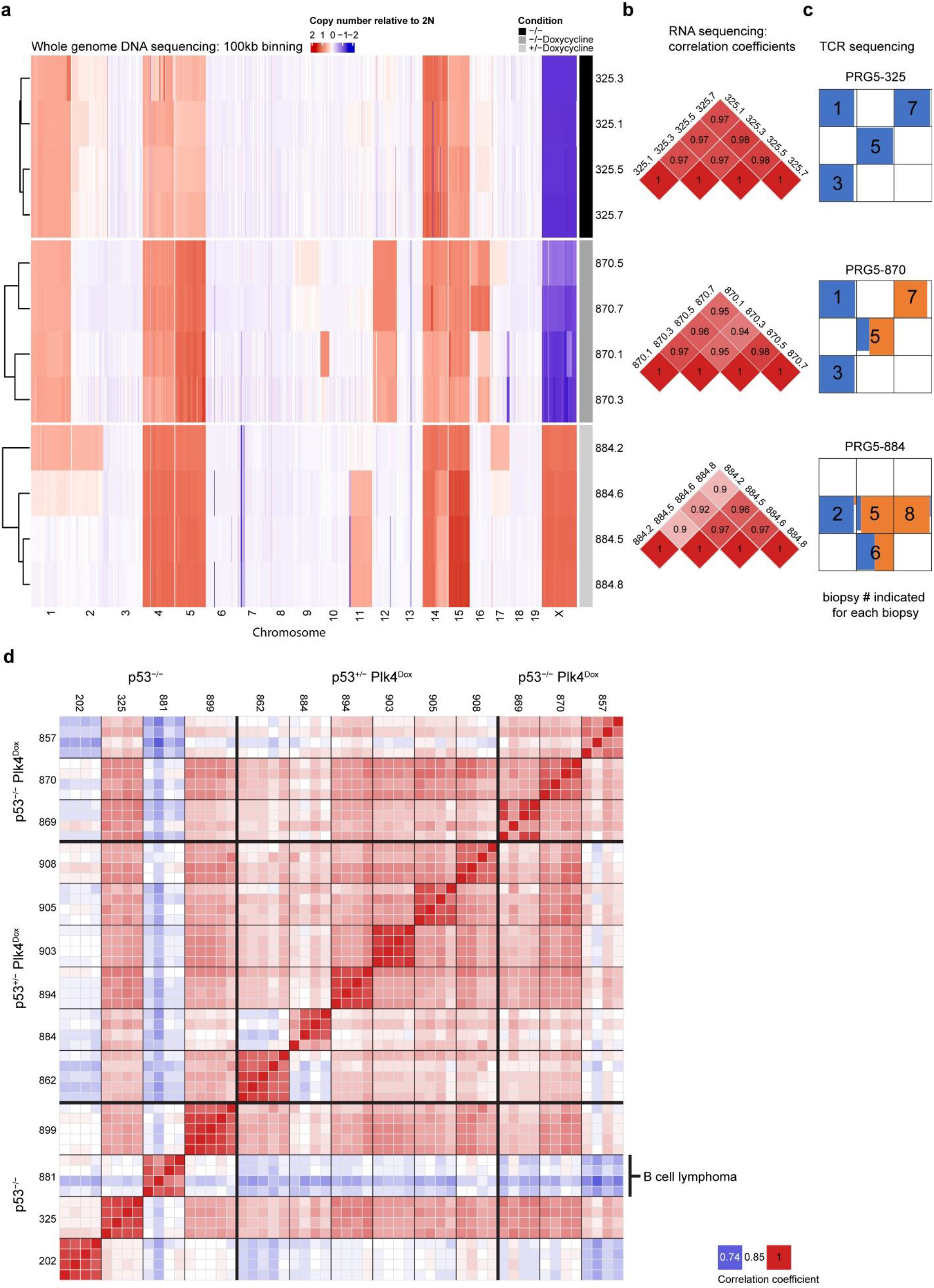
DNA and RNA levels in multifocal biopsies of thymic lymphomas. (**a**) Heatmap showing 100-kb DNA copy number changes in multiple regions taken from late tumors (>500mg) from non-induced p53-/- PRG5 mice (black), two-weeks doxycycline treated (at the age of 30 days) p53-/- PRG5 mice (dark grey) and p53+/- PRG5 mice (light grey) as determined using whole-genome sequencing. (**b, d**) Heatmaps showing pair-wised Pearson correlation coefficients among samples on TPM values as determined using RNA sequencing of the indicated PRG5 tumor biopsies. (**c**) Top ten T cell receptor frequencies (indicative of T cell clones) in thymic T cell lymphomas from PRG5 mice in multi-regional biopsies as determined using T cell receptor sequencing (these samples are also shown in Figure 3k and presented here for convenience).

**Supplementary Figure 11.**
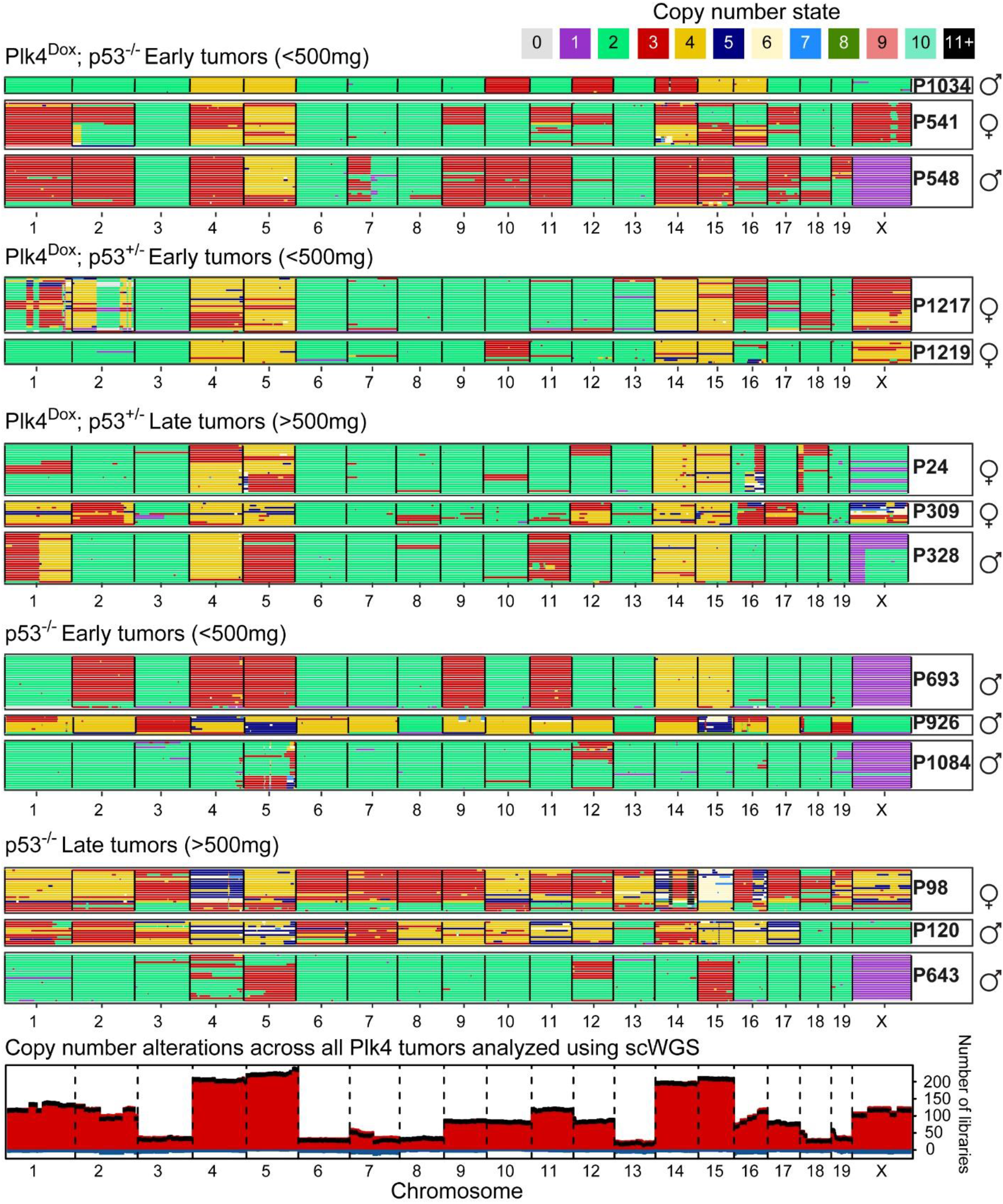
Single cell DNA sequencing of early and late PRG5 tumors. (**a**) Heatmaps showing DNA copy number using single cell whole-genome sequencing of Plk4 induced or control early and late PRG5 tumors with the indicated p53 backgrounds. Genomic position in order from chromosome 1 to X are in the x-axis and individual cells are in the y-axis. Colors indicate the copy number state as determined by AneuFinder. (**b**) Genome-wide overview of cumulative copy number (1 Mb bins) gains (red) and losses (blue) across all thymic lymphomas presented in panel **a**. Black line presents the net change; difference between number of libraries with a copy number gain and the number of libraries with a copy number loss.

**Supplementary Figure 12.**
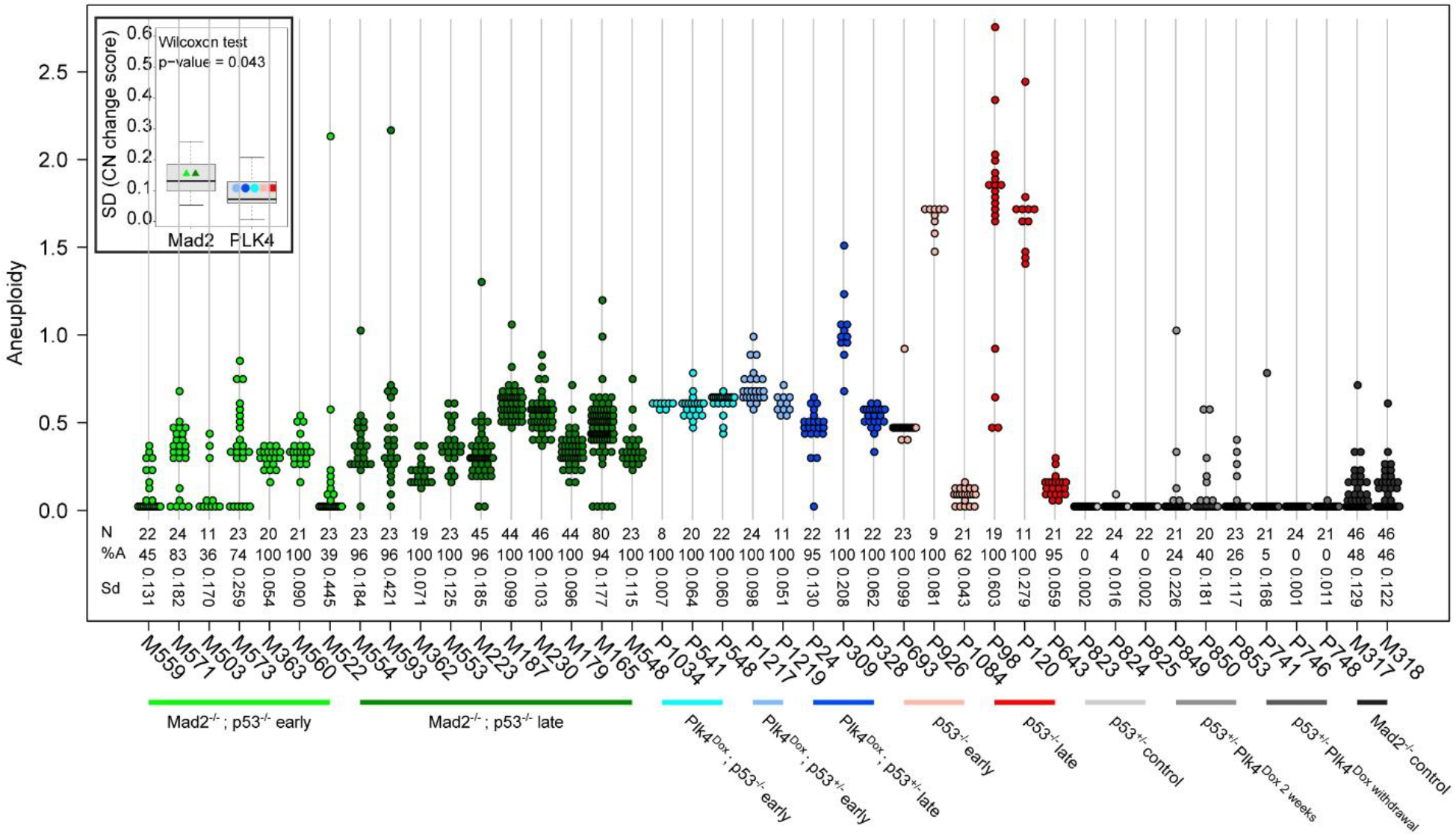
Differences between chronic and transient CIN in driving thymic lymphomas. Analysis of copy number changes of the entire Mad2 and PRG5 cohorts. The numbers below the dot plots indicate the total number of cells (N), the percentage of cells that has at least one whole chromosome gain or loss (%A) and the standard deviation of the copy number change scores of each sample (Sd). The insert boxplot shows the distribution of standard deviation values of the Mad2 and PLK4 samples. A Wilcoxon rank-sum test was used to determine if there is a significant difference between the SD values of the Mad2 and PLK4 samples.

**Supplementary Figure 13.**
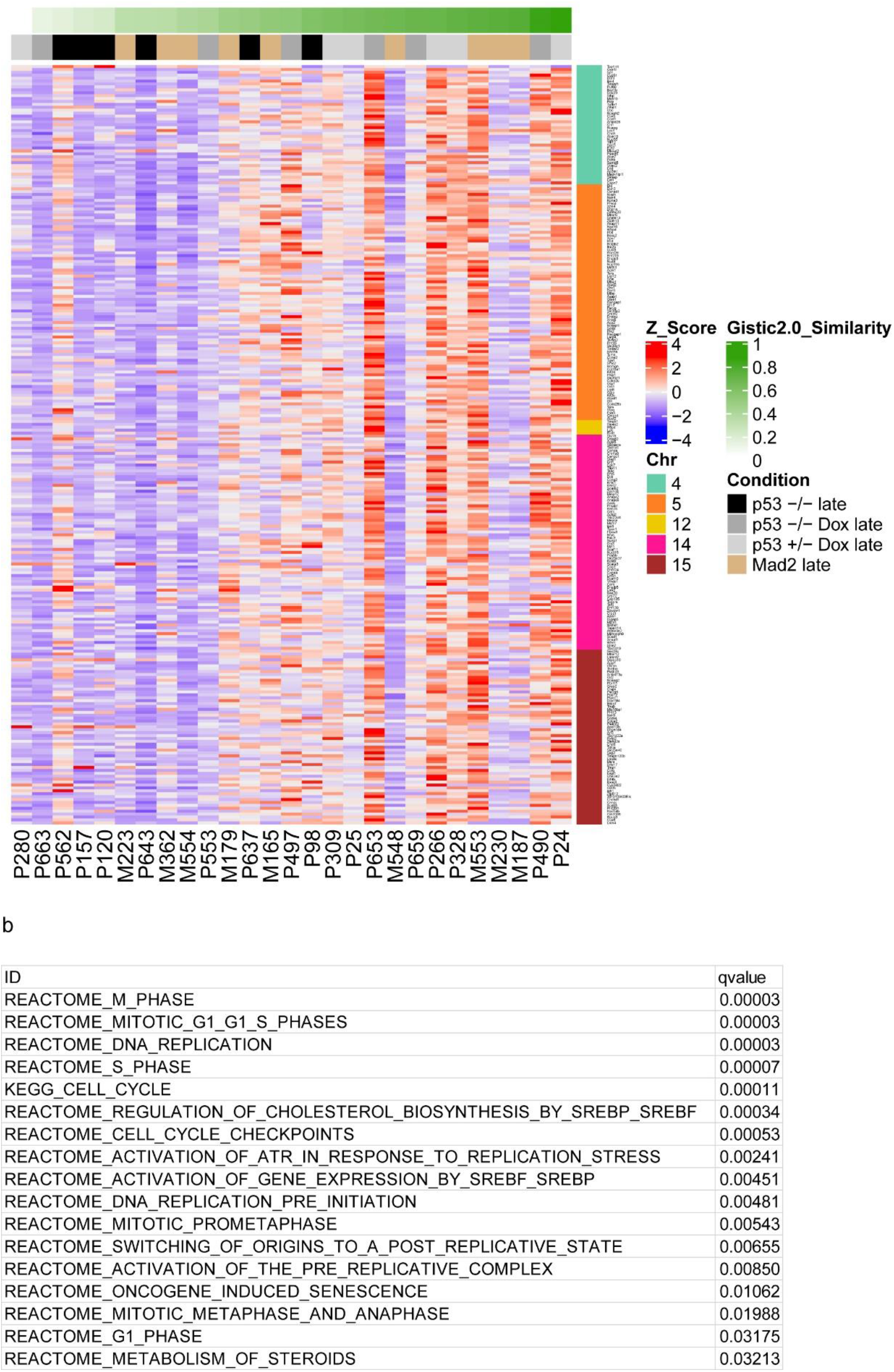
Acquisition of an aneuploidy specific profile reshapes expression of genes in cell cycle, replication, and stress networks. (**a**) Gene expression analysis from RNA sequencing preformed in correlation with the similarity score derived from comparison to the GISTIC 2.0 score (as shown in Figure 4) of terminal tumors from the indicated PRG5 and Mad2 mice. See Supplementary Table 5 for list of genes. (**b**) Significantly enriched pathways (identified using msigdb and R/Bioconductor package clusterProfiler) in PRG5 and Mad2 derived thymic lymphomas (as shown in panel a).

**Supplementary Figure 14.**
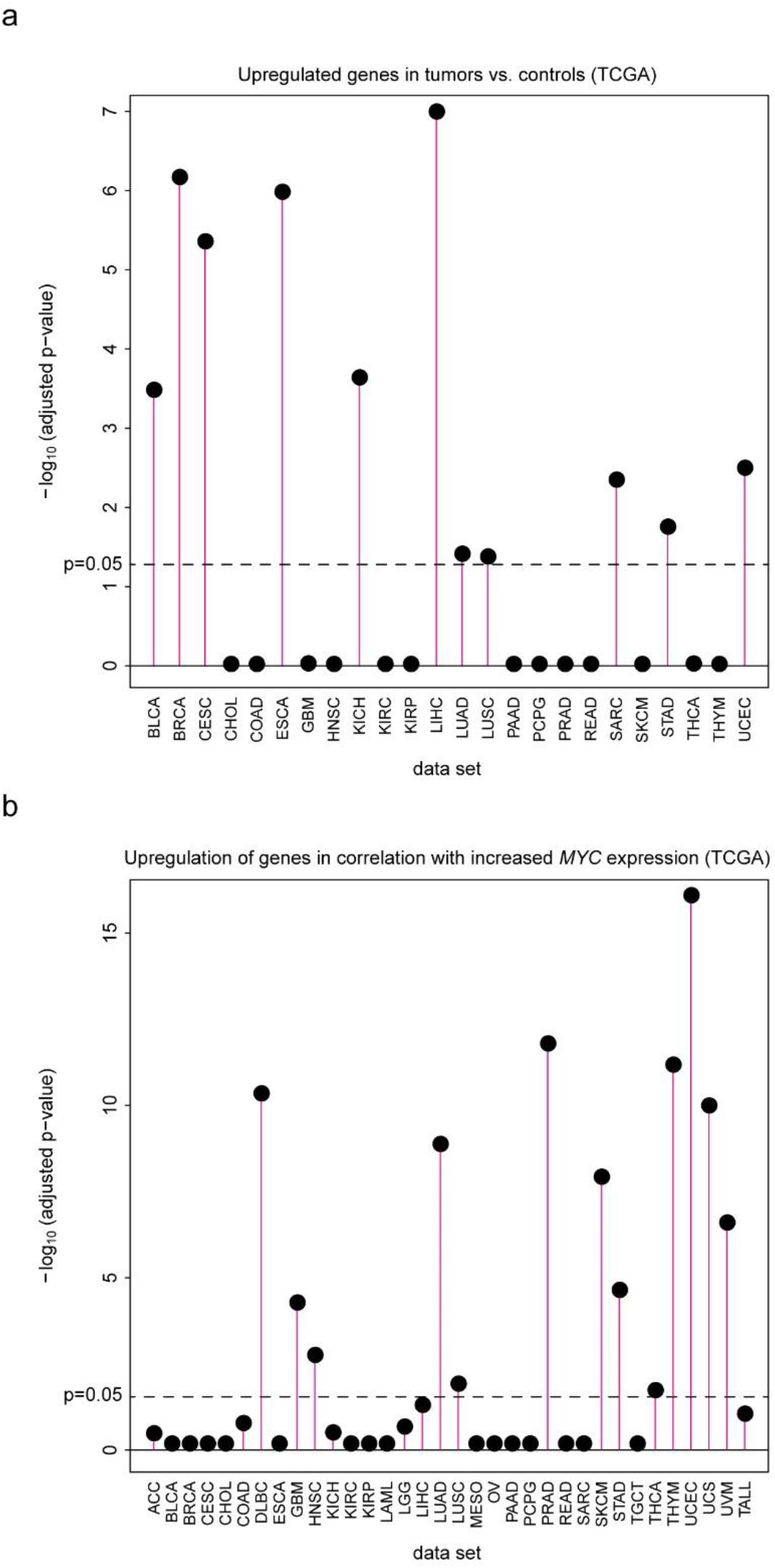
Multiple human cancers acquire the CIN induced gene expression profile, also in correlation with increasing MYC expression. (**a**) Enrichment of the CIN induced gene expression profile (Supplementary Table 5) in TCGA tumor cohorts gene lists of significantly upregulated genes relative to their non-tumor controls. (**b**) Enrichment of the CIN induced gene expression profile (Supplementary Table 5) in TCGA cohorts in correlation with *MYC* expression. A spearman correlation coefficient between the gene’s expression and MYC expression (among all samples), and the posterior probability that the gene is correlated with MYC was calculated. The ranked gene lists were then tested for enrichment with Bioconductor fgsea on the CIN induced gene expression profile (Supplementary Table 5).

## Supplementary Tables (see attached files)

**Supplementary Table 1 | Pathological analysis of tissue sections from PRG5 mice.**

**Supplementary Table 2 | Differentially expressed genes between thymic tumors and control thymuses as determined using RNA sequencing.**

**Supplementary Table 3 | Top 10 T cell receptor sequences sequences per sample as determined by T cell receptor sequencing**

**Supplementary Table 4 | Similarity to the GISTIC profile of PRG5 and Mad2 tumors (score ranges between 0-lowest and 1-highest).**

**Supplementary Table 5 | Gene expression analysis in correlation with the similarity index of each tumor in PRG5 mice (values represent Z scores)**

